# Serotype-Specific Detection of Non-Structural Protein 1 from Dengue Viruses by Surface-Enhanced Raman Spectroscopy: An Enhanced Precision Diagnosis

**DOI:** 10.64898/2026.06.13.731912

**Authors:** Monika Ghalawat, Virendra Kumar Meena, Atanu Basu, Pankaj Poddar

**Author notes:** Joint Corresponding Authors. **Corresponding Author Information** 1. Dr. Pankaj Poddar, Senior Principal Scientist, Physical & Materials Chemistry Division, CSIR- National Chemical Laboratory, Dr. Homi Bhabha Road, Pune-411008, Maharashtra, India.; 2. Dr. Virendra Kumar Meena, Scientist-D, Electron Microscopy & Histopathology Laboratory, ICMR-National Institute of Virology, Pune, 20-A, Dr. Ambedkar Road, Pune - 411 001, Maharashtra, India. Equal Contributors (Monika Ghalawat, Virendra Kumar Meena).

## Abstract

Dengue disease exhibits diverse clinical manifestations in patients when infected by its different serotypes. Early and accurate detection of dengue virus (DENV) infections, particularly distinguishing between serotypes is crucial for effective patient management and sporadic outbreak control. Surface-Enhanced Raman Spectroscopy (SERS) offers advantages of high sensitivity, rapid acquisition, rapid analysis, minimal sample and preparation requirements. In this study, we present a simple and reproducible approach for serotype-specific detection of non-structural protein 1 (NS1) utilizing SERS on an aluminium based substrate. Leveraging specific vibrational signatures of NS1 protein from DENV serotypes, we demonstrated the potential of SERS to discriminate between NS1 proteins across DENV serotypes and also the amino acid residue variations that exist among them from different biological samples. Study demonstrates the SERS based detection of NS1 in the current *in-vitro* setting has sensitivity and specificity comparable to ELISA assays with limit of detection (LOD) reaching to 1ng/mL. However, the application of nanomaterials-based SERS substrate has potential to further enhance the LOD enabling detection even at lower concentrations. This approach holds promise for advancing our capacity to rapidly diagnose serotypic DENV infection in samples, studying pathogenesis and improving strategies for disease management and control.

## INTRODUCTION

Non-structural protein 1 (NS1) of the DENV plays a pivotal role in viral replication, immune evasion, and pathogenesis of the disease, making it an attractive target for diagnostic assays and therapeutic interventions. Clinical spectrum ranges from acute febrile illness called dengue fever (DF) to severe life-threatening forms such as dengue hemorrhagic fever (DHF) and dengue shock syndrome (DSS) causing vascular leakage that leads to mortality. There is no specific therapy available for severe DENV infection till date, however clinical management of the patients primarily relies on fluid replacement therapy to preserve hemodynamic stability. Extensive genetic diversity among multiple circulating DENV serotypes in today’s time, led to the emergence of NS1 sequence variations, poses enormous challenges for conventional diagnostic techniques. This variability often leads to error in serotype identification, and contributes to delays in appropriate and timely clinical interventions.

Dengue diagnosis can be performed on various clinical samples such as serum, plasma, whole blood, and tissues from the liver, spleen, lymph nodes, lung, and brain collected from fatal cases. Conventionally available methods for dengue infection diagnosis are virus isolation, nucleic acid amplification assays, detection of antigens, and serological tests. Virus isolation may be employed in limited laboratory settings having necessary infrastructure and technical expertise. Virus isolation provides highly specific detection, however its extended reporting time (a week) limits its utility for early diagnosis. Nucleic acid detection using reverse transcriptase polymerase chain reaction (RT-PCR) requires longer procedure, time and expensive equipment with reagents. Test outcomes are highly dependent on personnel expertise, therefore high chance of manual error occurrence. Detection of NS1 antigen in serology-based tests are specific but its sensitivity varies widely and is less significant in secondary infection compared to the primary infection. Antigens and serological tests by ELISA also require certain facilities and take about a few days to produce results. The rapid diagnostic kits give results rapidly but have not been properly validated and have sensitivity issues as well. Therefore, rapid, sensitive, scalable, cost-effective and field-deployable diagnostic approaches are needed for early dengue diagnosis and reducing the global burden of the disease.

Raman spectroscopy is known to be a popular bioanalytical tool in order to detect and identify different biochemical species. It is also referred to as the fingerprint of molecules because of unique spectra that correspond to certain specific vibrational bands of a material. However, biologicals are weak Raman scattering materials with high fluorescence. The discovery of Surface Enhanced Raman Scattering (SERS) has tremendously increased the applicability of Raman spectroscopy. The rapid amplification of the electromagnetic field generated by the excited local surface plasmons on the noble metal nanostructures leads to the enhancement of the Raman signal of an adsorbed analyte molecule. It also provides the uniformity as well as the crystallinity of nanorods array having a high aspect ratio and localized surface plasmon resonance effect. Morphology and dimensions of SERS substrates are crucial parameters in order to have high value of enhancement. The specific reproducible surface morphology offers a high level of enhancement. A large variety of nanostructures have been found to manifest the SERS effect, including rough metallic surfaces by chemical etching, island films, aggregates of colloidal particles, nanorods, and nanowires fabricated by chemical and electrochemical methods or regular nanoparticle arrays prepared by nanosphere lithography or electron-beam lithography. However, many of these fabrication methods are either expensive, time-consuming, unable to reproduce the uniform morphology and maximum SERS enhancement. Therefore, SERS has been employed as a potential tool nowadays, for the label free detection of various bacteria and viruses. To date, there are only a few reports on SERS based DENV detection. Huh et al. demonstrated the detection of nucleic acid sequences of DENV-2 using a microfluidic SERS chip composed of electro-kinetically active microwells. Ngo et al. also detected the same employing plasmonic SERS active substrates. Paul et al. demonstrated DENV-2 detection using gold nanoparticles. However, in most of the SERS based detection studies, fluorescent dyes were labeled with target biomarkers. Therefore, a direct, rapid, field-deployable (on site diagnosis), user-friendly, reagent-free, and cost-effective diagnostic tool is the need of the hour. The presented technique meets most of the requirements in the form of an all-in-one set up which also provides the accurate results after matching with the stored database. In addition to that, it may offer early diagnosis which is of utmost importance in the disease treatment.

In this study, we propose utilization of SERS based detection of NS1 protein and their sequence variation among all four dengue virus serotypes, aiming to enhance diagnostic precision and accuracy. Our hypothesis is rooted in the premise that the unique spectral fingerprints obtained from NS1 proteins can aid to discriminate between serotypes and identify specific sequence variation associated with virulence and pathogenicity of DENV strains. Through comprehensive spectroscopic characterization and statistical analysis, we validated the sensitivity and specificity of SERS in distinguishing NS1 proteins from DENV serotypes and identifying their sequence variation. Successful implementation of SERS-based NS1 detection holds tremendous promise for transforming dengue virus diagnostics, particularly in resource-limited settings where conventional methods may be impractical or inaccessible. By providing rapid and accurate serotyping capabilities, this novel approach has the potential to streamline clinical decision-making, improve patient outcomes, and contribute to more effective disease control strategies on a global scale.

## EXPERIMENTAL SECTION

### Material Used

Commercial grade Aluminium foil (standard thickness: 0.011mm), Recombinant purified NS1 proteins (DENV-1 NS1, DENV-2 NS1, DENV-3 NS1 and DENV-4 NS1), Commercial grade human serum (MegaCult-C; Catalog No: 04807) from StemCell Technologies, Standard glass slide and Scotch tape.

### Instrumentation

An HR 800 Raman spectrophotometer (Jobin Yvon, HORIBA, France) was used for spectra acquisition. An achromatic Czerny–Turner type monochromator (800 mm focal-length) with silver-treated mirrors equipped with a Peltier cooled He–Ne laser light source (633 nm) operating at 20 mW was used as the illumination source. Emitted signals were captured using spectroscopy grade multi-channel CCD detectors (1024 × 256 pixels of 26 μm) under low dark current conditions (<2 × 10-3 pixels/s) with a 50x LD (Long distance) magnification. Silicon reference was used for the equipment calibration.

### Purification of Recombinant NS1 Proteins

Full-length recombinant NS1 proteins (DENV-1, DENV-2, DENV-3 and DENV-4) were purified using 6X His-tag complexed on Ni-NTA column chromatography after expressing the cDNA NS1 clone in pCMV-3 expression vector in HEK-293T cells. After purification, the NS1 proteins were concentrated using the dialysis method. Subsequent to purification of NS1 proteins, concentrations of NS1 protein from each serotype was obtained using BCA protein estimation assay. NS1 proteins of all the DENV serotypes were purified using a similar approach and cell culture protocols.

### SERS Spectra of Purified DENV NS1 Proteins

For acquiring the characteristic spectral signature peaks of NS1 proteins across DENV serotypes, SERS spectra measurements were performed of NS1 protein derived from each serotype. Prior to spectral acquisition, instrument calibration was carried out using a silicon reference standard. DENV NS1 protein samples were diluted in nuclease- and protease-free double-distilled autoclaved water to a final concentration of 0.5 µg/µL. Two sequential drops (1.0µL each) of the diluted NS1 protein solution were deposited onto an aluminum foil substrate supported on a glass slide and allowed to air-dry at laboratory temperature (∼25 °C) for 10–15 min to form a uniform dried film. Raman spectra were recorded using an acquisition time of 5 s and 10 s. Spectra were collected over the wavenumber range of 200–3000 cm⁻¹ with a spectral resolution of ±1 cm⁻¹. SERS spectra of NS1 proteins from all four dengue virus serotypes were acquired using the same experimental procedure to ensure uniformity and comparability of spectral data. All measurements were performed at room temperature (∼25 °C) under identical instrumental conditions.

### Detection of DENV NS1 Proteins in Serum

#### Spectral Uniformity and Reproducibility of NS1

The SERS spectra were acquired from three different locations on the same sample, and the resulting spectra were carefully compared in order to investigate the spot-to-spot consistency of the spectra in NS1 samples. To confirm sample-to-sample reproducibility, additional independent measurements were performed on DENV-3 NS1 and DENV-4 NS1.

#### Human Serum

To examine the applicability of the proposed method under more complex biological conditions, HS was used as the sample matrix. Primarily, pure HS and NS1 solutions were tested separately; after that, HS–NS1 mixtures were prepared in equal volume ratio (1:1). The obtained HS–NS1 samples were drop-cast on the Al foil substrate and examined.

To select an optimal HS dilution for detecting NS1 protein in complex biological conditions, HS was serially diluted (10^-1^–10^-4^) in nuclease- and protease-free double-distilled autoclaved water. SERS spectra were acquired for all dilutions to investigate their effect on the quality of the obtained SERS spectra.

Furthermore, DENV-2 NS1 was mixed with all four diluted HSs to make the final concentrations 1 × 10^5^ and 1 × 10^4^ ng/mL, and SERS spectra were acquired. Finally, an ideal HS dilution for obtaining the finest NS1 signals was chosen based on the quality and clarity of SERS spectra. Additionally, the reproducibility was determined for the selected HS dilution through three independent measurements with an NS1 concentration of 1 × 10^4^ ng/mL.

#### Various Concentrations of NS1 in Diluted Human Serum

Once the HS dilution has been optimized, the effects of NS1 concentrations in the presence of diluted HS are investigated for the lowest detection limit. NS1 proteins from all four serotypes (DENV-1 to DENV-4) were prepared to make the final concentrations of 5×10^5^, 5×10^4^, 5×10^3^, 500, 50, and 5 ng/mL in diluted HS. These solutions of NS1 proteins were drop-cast on the Al foil substrate for the detection of NS1 proteins in the mixture.

#### MATLAB Pre-processing

The Raman spectra were left unprocessed and shown in their original form for general display purposes. Nevertheless, only deconvolution studies were preprocessed. The spectra were loaded into Origin 9 first, and the region of interest (320 to 2000 cm^−1^) was selected. The spectra were normalized over this range to facilitate comparison. Subregions were then selected for fitting in MATLAB after zooming the regions.

To enhance the quality of a raw spectrum and allow for reliable quantification or analysis, pre-processing of the Raman spectra was carried out using our built MATLAB program. The smoothing of the raw spectrum (intensity versus Raman shift) was completed by applying the Savitzky–Golay filter algorithm (second-order polynomial, 11-point). ALS method was used to perform the baseline correction operation. By this mechanism, we can obtain an estimate of the background signal that penalizes deviation from the baseline. In this instance, λ was predetermined to be 10⁵, and p was set to 0.001 in order to avoid distortion of signal peaks while capturing the background of the fluorescence signal. To get the final spectral data, the baseline estimation was taken away from the smoothed spectral data, leaving background-free spectra ready for further analysis (peak fitting). The resultant spectra then went through Gaussian peak deconvolution using our MATLAB program. For the initial peak positions, we picked out clearly defined peaks and shoulder features in the spectra. During the fit, we made sure each peak’s full width at half maximum (FWHM) was kept between 10 and 25 cm^-1^ to make physical sense. We judged the fit quality using the coefficient of determination (R²). Good fits had an R² of at least 0.98. Some spectra had slightly lower R² values between 0.96 and 0.97 but still matched up pretty well with the experimental data.

We kept the number of fitted peaks and the fitting strategy constant for all spectra to make sure we had reliable quantitative comparisons, especially for samples that only differed in NS1 concentration. This reduced fitting variations and ensured any changes in peak area were because of real spectral differences and not math artifacts from different models. Next, Gaussian deconvolution was used for quantitative analysis based on peak areas. After that, we plotted concentration-dependent trends to see the spectral changes for various NS1 levels and to explore how peak position and area depended on NS1 concentration.

## RESULTS AND DISCUSSION

This study focused on using Raman spectroscopy to spot the four types of dengue viruses, each identified by their non-structural protein 1 (NS1). Although Raman spectroscopy is expert at giving precise molecular info, it struggles with biological samples because they don’t scatter light efficiently. Researchers overcame this problem by boosting the technique with surface-enhanced Raman spectroscopy (SERS). SERS intensifies the signal using localized surface plasmon resonance. In this study, we even figured out a low-cost way to do this by using ordinary aluminum foil as a SERS substrate.

### Substrate of Aluminium Foil

We used available commercial aluminium foil as our substrate and made sure to use the bright side to boost surface reflectivity. Figure 1 shows the difference between the shiny and non-shiny sides of the foil. With the FESEM and AFM imaging (Figure 1A1-C1 and A2-C2) in different orientations, we found that the non-shiny side was rougher. EDAX maps for both sides are shown in Figure 1D1-D2, indicating the presence of pure aluminium. Due to this high roughness and lower reflectance, the non-shiny side was inappropriate for SERS; therefore, the shiny side was used in this study as a SERS substrate. It worked better because it was smoother and had better reflectance, perfect for those plasmonic amplifications. Additionally, using commercially available Al foil is even better than synthesized nanoparticle substrates, as its large-scale production keeps the surface uniform and consistent from batch to batch. This lowers variations between substrates, cuts down on measurement differences, and makes SERS analyses more reliable. In conclusion, Al foil works as a great substrate for affordable and scalable diagnostics.

**Figure 1.**
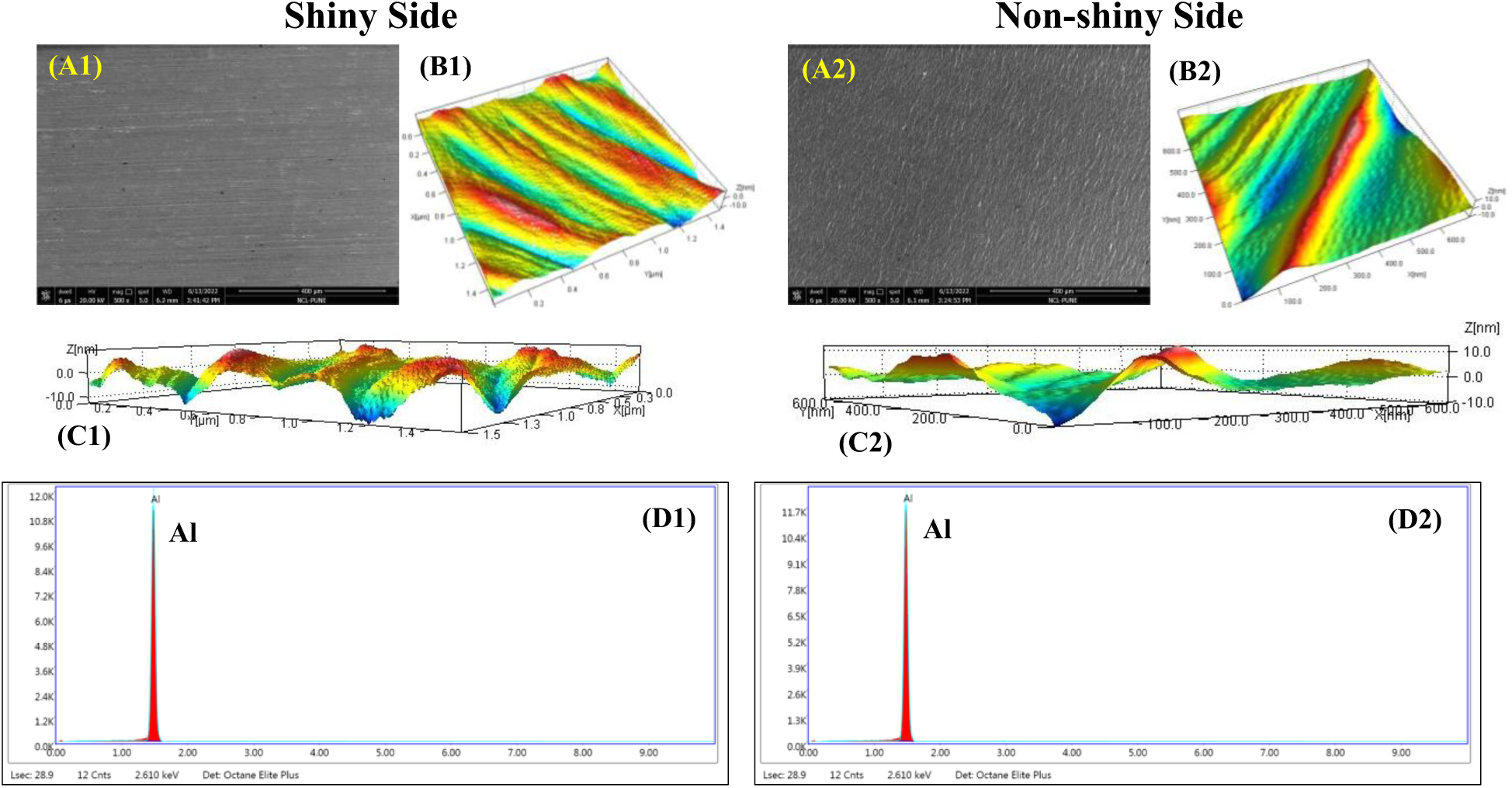
SERS substrate characterization of aluminum foil: (A1–D1) shiny side and (A2–D2) non-shiny side. (A1, A2) FESEM and AFM images—(B1, B2) vertical and (C1, C2) horizontal surface profiles for comparing the morphology assessment. (D1, D2) EDAX spectra confirming elemental composition on both sides.

### SERS Detection of Different Serotypes of DENV

Figure 2 shows the full-length NS1 protein sequences that vary among four dengue viruses. The amino acids are represented by single-letter codes. The conserved residues are marked with colors, whereas non-conserved ones are left white. A zoomed-in view of these differences is shown in Figure S1 (only non-conserved residues). These sequence variations cause distinct vibrational spectra when analyzed through SERS. The NS1 protein from all four serotypes is analyzed by dropping the samples onto shiny aluminum foil (2 µL, repeated twice) and letting them air dry for ∼25 ± 5 minutes. SERS spectra were recorded from 200 to 3000 cm⁻¹, with a resolution of ±1 cm⁻¹.

**Figure 2.**
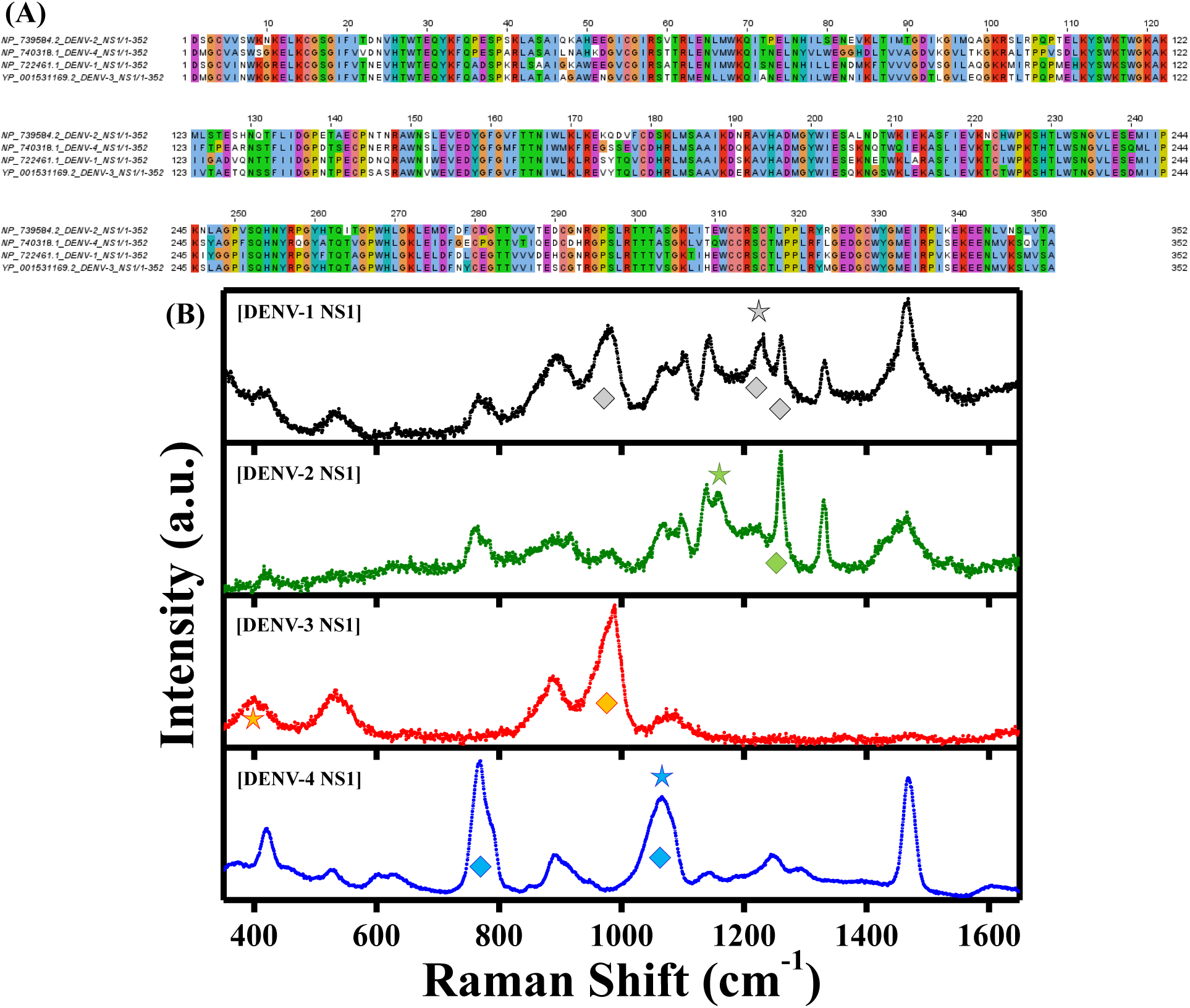
(A) Multiple sequence alignment of full-length NS1 proteins from different DENV serotypes, highlighting amino acid variations. Residues are represented using single-letter codes, with conserved residues color-coded and non-conserved residues shown in white. (B) SERS spectra of purified recombinant NS1 proteins from DENV serotypes (1.0 µg), exhibiting distinct spectral fingerprints. Star symbols denote serotype-specific signature peaks characteristic of each NS1 protein, while diamond symbols indicate the primary serotype-specific peaks that are critical for subsequent sample analysis.

The control experiments show that the signals we got are indeed from SERS-based tests. When we put NS1 on a glass slide, we only got a weak spectrum with no clear peaks (see Figure S2A). Also, when we checked an empty aluminum substrate, there were no peaks, excluding the contribution from the substrate (Figure S2B). Therefore, it is appropriate to conclude that the obtained spectra are coming from the SERS effect of NS1 proteins sticking to aluminum foil.

We examine the spectral uniformity over all four serotypes (spot-to-spot consistency). Spectra are taken from several spots on the same sample, and they show similar peak positions and intensity, meaning the SERS response is nearly consistent over the sample. Figure S3 illustrates that the peaks are nearly identical with very little variation, proving the substrate’s homogeneity. In order to determine the reproducibility of the spectra, independent measurements were performed. These spectra for DENV-3 NS1 and DENV-4 NS1, even from different positions (see Figure S4), confirm high reliability and reproducibility. As the spectra were recorded from various spots on the same sample. After comparing them and verifying the spectral profile, the spectra with the high signal-to-noise ratios were selected for further serotype comparison.

SERS spectra for NS1 proteins from the four serotypes of the dengue virus are different (Figure 2B). For better visualization, the spectra are color-coded: DENV-1 is black, DENV-2 is green, DENV-3 is red, and DENV-4 is blue. Specific peaks are observed for each serotype, marked with stars, showing a unique fingerprint for each serotype. In addition, each serotype showed the high-intensity bands marked with diamond symbols in Figure 2B; their importance will be discussed further in the following parts. Notably, rather than a single feature, it is actually the collective pattern of bands that enables the serotype recognition.

The serotype spectra with the position of diagnostic bands are provided in Figure S5, which together highlight 20 Raman bands spread across all the serotypes. Some of these bands are unique to specific serotypes, making it easier to tell them apart. DENV-1 NS1 shows up at 1230 cm⁻¹, DENV-2 NS1 at 1159 cm⁻¹, DENV-3 NS1 at 399 cm⁻¹, and DENV-4 NS1 at 1065 cm⁻¹. Table S1 tabulated these positions along with tentative assignments for both shared and unique vibrational modes. Additionally, the color intensity in Table S1 corresponds to the relative band intensity. This process shows that SERS is a powerful technique for spotting the NS1.

Comparing SERS spectra across four DENV serotypes shows an interesting mix of shared vibrational fingerprints and unique changes. We identified three shared bands (present in all serotypes) at 533, 887, and 1468 cm^-1^ that mark key parts of the NS1 protein framework. The 533 cm^-1^ peak is due to disulfide bridges stretching, the gauche-gauche-gauche (g-g-g) conformation of cystine residues. It is strongly present in DENV-1, DENV-3, and DENV-4 but only faintly appears in DENV-2. That weaker signal means DENV-2 is either more flexible or folds differently. Another band at 1468 cm^-1^ linked to CH2 and CH3 deformation in side chains holds up well in DENV-1, DENV-2, and DENV-4; it noticeably fades in DENV-3, reflecting variations in the hydrophobic packing within the lipid-binding “wing” domains of the hexameric NS1 structure. Crucially, the band at 887 cm^-1^, which is linked to skeletal C-C stretching and tryptophan ring breathing, stays consistently strong in all four serotypes. This feature makes it really useful, as it is a stable and universal spectral signature peak for NS1 protein in all serotypes. We can rely on this 887 cm^-1^ signal to accurately detect the NS1 protein in complicated samples that is for “positive/negative” diagnostic screening of dengue virus. When they spot this band, along with the 533 cm^-1^ and 1468 cm^-1^ bands, they know for sure it is a “positive” for the NS1 protein. After that, more detailed analysis in the high-resolution fingerprint regions helps in determining the serotyping. In conclusion, we first use mainly the 887 cm^-1^ peak signal to check for the NS1 protein, then inspect more closely to identify the specific serotype. It is a rigorous, fast, and reliable technique to meet the high standards needed for diagnosing dengue.

In NS1 spectra, distinct serotype-specific marker bands are seen, giving clear spectral differentiation. Moving on to the non-conserved peaks in serotypes: there is the low-wavenumber band at ∼399 cm⁻¹ found only in DENV-3, which is linked to skeletal deformations. It might be amplified by specific molecules–substrate interactions during SERS conditions. Next up is the peak at ∼420 cm⁻¹; it points to out-of-plane bending and C–S/S–S vibrations. These show us cysteine residues and disulfide linkages. Over in the ∼601 cm⁻¹ range, we get ring breathing modes related to phenylalanine. Meanwhile, the ∼767 cm⁻¹ mark represents indole vibrations from tryptophan. There are also extra features around ∼916 and ∼989 cm⁻¹ that come from protein backbone vibrations and aromatic amino acid residues. Another key spot is at ∼1065 cm⁻¹ for DENV-4, corresponding to C–N stretching and aliphatic C–C vibrations, suggesting differences in backbone and side-chain shapes. Lastly, the region ∼1100–1300 cm⁻¹ includes contributions from C–C, C–N stretching, and amide III mode contributions—from 1103, 1139, 1244, and 1260 cm⁻¹—that really give insight into protein conformation and amino acid compositions.

Higher wavenumber bands around ∼1292, ∼1330, and ∼1468 cm⁻¹ show CH₂/CH₃ deformation and twisting modes, highlighting the protein composition and side-chain dynamics. The ∼1230 cm⁻¹ band (only present in DENV-1) lies in the amide III region and points to N–H bending and C–N stretching, sensitive to the secondary protein structure. As for the ∼1159 cm⁻¹ peak, only present in DENV-2, it comes from C–C and C–N stretching, and from differences in side-chain compositions and local chemical environment. These specific peaks show up distinctly in their respective types, making them dependable markers.

Human serum (HS) offers a real-world, complex environment for examining the efficiency of the proposed SERS detection method. Figure S6 illustrates the spectra of pure HS, pure NS1, and a mix of both in a 1:1 ratio for all four Dengue virus serotypes. The pure HS spectrum acts as a reference to reveal the contribution from serum components.

Among the four serotypes, DENV-2 NS1 shows the strongest spectral response when mixed with pure HS. As emphasized by the arrow in the shaded region of Figure S6B, the spectra keep two clear NS1 bands visible even in the complex HS mix, demonstrating that the platform can easily spot NS1 against the complex background also. In comparison, DENV-1 and DENV-4 show some changes too after being mixed with HS, shown in the shaded region of Figures S6A and S6D. However, these changes are not as prominent as DENV-2. The clearly visible changes in the DENV-2 NS1:HS mixture as compared to pure HS spectra make it easier to examine the presence of NS1, making NS1 detection simpler. In conclusion, the NS1 spectral signatures stay recognizable in real biological situations. So, the aluminum foil-based SERS system shows the potential for directly finding dengue NS1 in serum samples.

In view of its high-sensitivity spectral response, as mentioned above, the DENV-2 NS1 was chosen as a specimen serotype in order to determine the optimal circumstances for NS1 detection in the presence of HS. Figure S7A illustrates the SERS spectra of pure HS and its dilutions from 10^-1^ to 10^-4^; characteristic serum bands are clearly visible up to 10⁻² dilution. Whereas the signal becomes noisier with further dilutions, making peak identification difficult. Next, to determine the optimal HS concentration for NS1 detection, we mixed DENV-2 NS1 at concentrations of concentrations 1 × 10^5^ and 1 × 10^4^ ng/mL with HS diluted from 10⁻¹ to 10⁻⁴, shown in Figures S7B and S7C. We also included spectra of pure HS and pure DENV-2 NS1 for comparison.

The comparison of the spectra shows that the distinctive NS1 band at 1260 cm^-1^ stands out best in an HS dilution of 10^-2^, as clearly highlighted in the shaded region. The NS1 signal gets covered up at higher serum concentrations, and at lower ones, the quality of the overall spectrum drops due to weaker signal intensity. To confirm, three separate tests were done using concentrations 1 × 10^4^ ng/mL DENV-2 NS1 in HS diluted to 10^-2^ (Figure S7D). Each test showed the 1260 cm^-1^ band consistently. Given these reliable findings, 10⁻² diluted HS was picked as the ideal condition for detecting NS1. It works great because it cuts down on serum interference while keeping the spectral quality up. Therefore, this specific dilution became standard for all further serum-based analysis.

After optimizing the HS dilution, the NS1 proteins of all four DENV serotypes were mixed with a 10^-2^ diluted HS at different concentrations from 5 × 10^5^ to 5 ng/mL to observe the effect of NS1 concentration on spectra and whether the unique signatures of each serotype stayed detectable (Figure S8). For each serotype, specific regions in the spectra really stand out. In DENV-1 NS1(Figure S8A), the most useful region for diagnosis is shaded in dark gray, with some extra helpful features in light gray. For DENV-2 (Figure S8B) and DENV-3 (Figure S8C), the special markers are in the dark green and dark orange regions, respectively. Last, for DENV-4 (Figure S8D), the main diagnosis happens in the dark blue area, with additional discriminatory information in the lighter blue region. The constructive peak area of approximately 887 cm^-1^ in all serotypes is indicated by the yellow-shaded region in Figure S8A-D.

At high NS1 concentrations, like 5 × 10^5^ ng/mL, you see contributions from both NS1 and HS components with NS1 dominance, which gives clear spectral protein characteristics. However, as NS1 concentration drops, serum bands start taking over. However, over the whole concentration range examined, the diagnostic areas indicated in Figure S8 maintain detectable spectral differences in comparison to the HS control. The characteristic spectral patterns last all the way down to as low as 5 ng/mL. This shows that even in the presence of significant matrix interference, serotype-specific spectral information can still be identified.

For example, when there is no NS1, a pure HS doublet appears as shown in the shade region in between 1200 and 1300 cm^-1^ (Figure S8A-B). However, in the presence of the smallest concentration of NS1 (5 ng/mL), that doublet changes shape—peaks get broader, and lines get altered. This observation indicates the formation of overlapping NS1-associated vibrational modes that disturb the native serum spectral profile. For DENV-3 and DENV-4 at lower concentrations, they mainly show changes in the intensity ratios between neighboring bands in raw data, not full peak mergers. This means the SERS platform stays really sensitive, spotting those shifts even at its lowest test levels.

The way the spectra evolve based on different concentrations shows we can spot NS1 not only by looking for unique marker bands, but also by tracking changes in peak shape, relative intensity, and band broadening in specific spectral windows. These features remain visible even at really low concentrations, stressing the Al foil-based SERS platform’s potential for detecting dengue NS1 proteins in complicated serum samples. Therefore, to further elucidate the spectral changes at different NS1 concentrations—where at lower concentrations their bands might overlap with those of serum—we used peak deconvolution analysis. This method enabled the hidden details and gave us a better understanding of the contributions made by NS1 and serum constituents.

### Resolving Overlapping Serum and NS1 Vibrational Bands through Peak Deconvolution

Although the raw spectra showed clear concentration-dependent spectral changes, we couldn’t right away figure out the much clearer difference due to overlap between serum and NS1-derived vibrational bands. Therefore, peak deconvolution was performed in the selected regions using Gaussian fitting, which provided the overlaps and helped us find NS1-specific modes hidden by the serum. As a result, the mechanistic explanation is achieved for the peak broadening, doublet merging, and intensity redistribution when NS1 was added.

As discussed in the above sections, the ∼887 cm^-1^ band is consistently observed in all four DENV serotypes, making it a great serotype-independent marker for the NS1 protein. It appears reliably in both pure NS1 samples and those with serum too, which means it is a prime band for diagnosing NS1, no matter the specific infecting serotype. Due to this consistent spectral marking, there is a strong basis for using it in simple positive or negative screens for NS1. Given that, we picked the 810–915 cm^-1^ region for detailed peak deconvolution analysis to examine more closely NS1-related spectral modification in HS. Figures 3A- D show the deconvoluted spectra for DENV-1 to DENV-4. These include the pure HS spectra (Figure 3A1-D1), a mixture of 10^-2^ diluted HS with the two lowest NS1 concentrations (Figure 3A2-D3), along with their respective pure NS1 versions (Figure 3A4-D4). The objective is to determine whether the NS1 signals are still detectable at the lowest NS1 concentration in the presence of HS (10^−2^).

**Figure 3.**
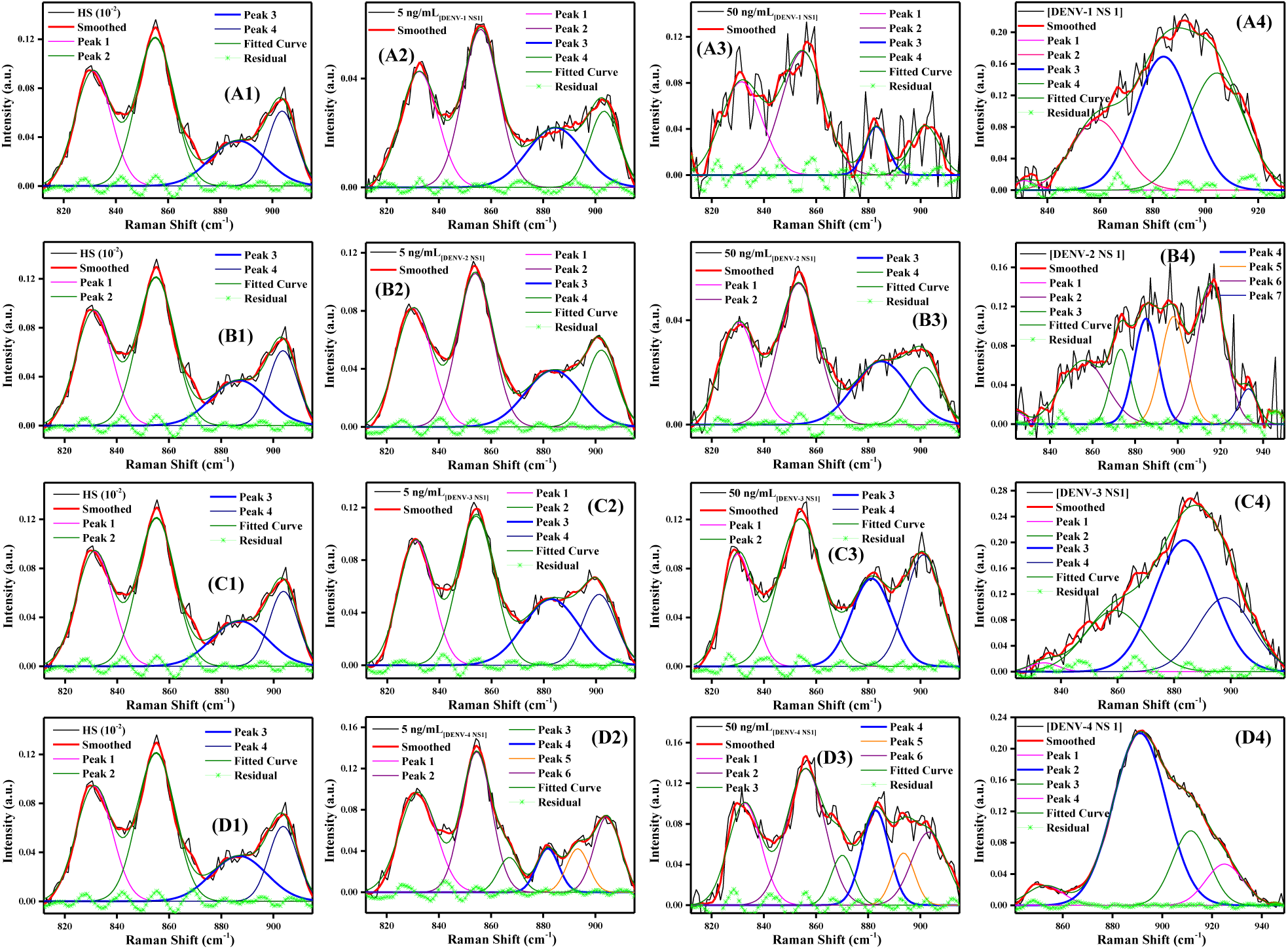
Deconvolution of Raman spectra: (A1–A4) DENV-1 NS1, (B1–B4) DENV-2 NS1, (C1–C4) DENV-3 NS1, and (D1–D4) DENV-4 NS1. Series 1–4 represent the diluted HS (10⁻²), mixtures of corresponding NS1 (5 and 50 ng/mL) with HS (10⁻²), and pure serotype-specific NS1, respectively.

The deconvolved spectra show that the NS1-containing samples differ from the HS controls. Serum spectra show clear peaks, and adding NS1 brings new vibrational components and changes the strength of the existing bands. In Figures 3A1–D1, when we look closely at the HS control, its deconvoluted spectrum fits four Gaussian components. Out of these, Peak 3 at around 887 cm^-1^ is really small. We picked this peak as it is present in all pure DENV NS1 serotypes, making it a perfect point for more checks. When NS1 is added at the lowest concentration of 5 ng/mL (Figures 3A2–D2), we notice changes near the 887 cm^-1^ band. These changes make it easier to tell which samples have NS1 at a 5 ng/mL concentration also. For DENV-1 and DENV-2, the related part moves to around 884 cm^-1^ and gets more intense compared to the serum control. Similarly, for DENV-3, the peak shows up between ∼881 cm^-1^ with increased signal strength too. The deconvolution data matches the raw spectra shown in Figures 3A2–C2, clearly demonstrating higher intensity in that region. With DENV-4 (see Figure 3D2), a new component pops up at ∼882 cm^-1^, which wasn’t there in the serum sample. As these shifts last through all serotypes too, this proves that this part of the spectrum is good for spotting dengue NS1 in general.

Further increasing the NS1 concentration to 50 ng/mL (Figures 3A3–D3) makes this spectral feature stand out even more for all serotypes. The way the peak evolves, shown by the blue peak, means the changes observed are due to NS1 and not any random serum noise. Certainly, to confirm the finding, the pure NS1 spectra are also deconvoluted using the same process (Figures 3A4–D4). The peaks are prominently observed ∼884 to 891 cm^-1^ in all serotypes, proving it belongs to NS1. In conclusion, the band ∼887 cm⁻¹ appears in all four dengue serotypes, with the lowest NS1 concentration also. It is consistently present in both pure and mixed samples, strengthening the idea that it is an NS1 marker. This 880–890 cm⁻¹ area can therefore reliably distinguish NS1-positive from NS1-negative samples before any further tests, making it really useful for quicker diagnoses. Therefore, the proposed method has the ability to detect a conserved NS1-related signal before determining the serotype. So, we first do universal NS1 testing, then use typical marker bands for specific virus types, as discussed earlier.

To identify each DENV serotype, the peak deconvolution is performed in the diagnostic spectral regions unique to them. Starting with DENV-1 NS1, we focused on the region around 960 cm⁻¹, shown in the light gray shaded part of Figure S8A. The deconvoluted spectra from this region are shown in Figure S9. In Figure S9A, the pure DENV-1 NS1 spectrum fits with three Gaussian components. The broad feature around 960 cm⁻¹ (Peak 1, blue curve) stands out as the key indicator. When you mix DENV-1 NS1 with 10⁻² diluted HS, the spectrum changes because of both components. At the highest concentration tested (5 × 10^5^ ng/mL), the NS1 parts still dominate, with the 960 cm⁻¹ band easily noticeable. As the NS1 concentration goes down, the serum bands get more prominent, making the NS1 signal weaker, as shown in Figure S9. However, even at the very lowest tested concentration (5 ng/mL), both the raw spectrum and the analysis show the 960 cm⁻¹ feature dominantly compared to pure HS (Figure S9G-H). This proves it is a great marker for DENV-1 NS1. To examine this band behavior at different concentrations, we observe the peak position and integrated peak area – these are summed up in Figures S9I and S9IJ. As the NS1 concentration goes down, the peak position shifts to higher wavenumbers. This indicates changes in the vibrational environment that depend on the concentration of NS1. At the same time, when NS1 concentration drops, the integrated peak area gets smaller too, matching the idea of NS1’s vibrational modes contributing less in the serum matrix. The ∼960 cm^-1^ peak is only one aspect of this region. Throughout the entire concentration series, there is a second feature at about 985-990 cm^-1^, which is really the strongest band in the spectrum of pure DENV-1 NS1. As the concentrations decrease, its intensity also decreases but does not completely disappear, as shown in Figure S9G, peak 5 at the lowest concentration. The persistence of both of these spectral features demonstrates the specificity of the DENV-1 NS1 fingerprint. This suggests that an accurate serotype can be identified even in situations with significant serum interference.

The second DENV-1 NS1-specific diagnostic region, shown by the dark gray shading in Figure S8A, was further examined using peak deconvolution—as seen in Figure S10. Figure S11 displays an enlarged representation of the key spectral features. In this region, two bands at around 1230 and 1260 cm^-1^ were identified as the main spectral markers for DENV-1 NS1. To accurately fit the data and reduce baseline artifacts, we did deconvolution over a wider spectral range that included all nearby vibrational features (Figure S10). This method helped us reliably separate the overlapping bands and avoid math errors from fitting a narrow spectrum with multiple unresolved parts in this region. The close-up views in Figure S11 show the key diagnostic peaks around ∼1230 and ∼1260 cm^-1^. The deconvoluted spectrum of pure DENV-1 NS1, shown in Figures S10A and S11A, clearly shows well-defined bands at around 1230 and 1260 cm^-1^, confirming they’re intrinsically part of the NS1 protein. When 10⁻² diluted HS is added, the spectral profile changes as serum vibrations start overlapping with the NS1 bands. At the highest NS1 concentration of 5 × 10^5^ ng/mL (Figure S11B), the 1260 cm^-1^ band (peak 7, royal blue curve) stays very noticeable, while the 1230 cm^-1^ band (peak 5, purple curve) loses some intensity. As the NS1 concentration drops, the intensities of both bands keep fading. Nevertheless, the fitted components can be spotted in all concentrations, even down to 5 ng/mL. A key observation is that both bands are not present in the HS control spectrum. Consequently, DENV-1 NS1 must be the cause of their presence in the NS1 and HS mixture samples, making them a very specific marker for detecting DENV-1. Figures S11I-L summarize the integrated peak position and peak areas. With concentration, the peak position varies minimally. Nevertheless, the peak area of both peaks decreases with decreasing NS1 concentration, supporting the contribution of NS1’s vibrational modes. This suggests that variations in the vibrational environment are dependent on the NS1 concentration. Therefore, the deconvolution analysis shows the ∼960, ∼1230, and ∼1260 cm⁻¹ bands work great as a fingerprint for DENV-1 NS1, even at the lowest concentration.

The DENV-2-specific region, marked by the dark green area in Figure S8B, was further analyzed using Gaussian peak deconvolution, shown in Figure S12 and magnified views in Figure S13. This spectral window is the same as the second one used for DENV-1. Even though the distinct DENV-2 marker around 1159 cm^-1^ showed up clearly in the pure NS1 spectrum (Figure 2), its close proximity to some strong serum bands made direct viewing tough in the HS-containing samples. These nearby bands caused big overlaps. Therefore, another prominent peak was used for deconvolution to search for other spectral signs that still work in the serum mix.

Similar to the DENV-1 NS1 example, deconvolution was carried out over a broad range. The deconvoluted spectrum of pure DENV-2 NS1 (in Figures S12A and 13A) shows a big band right around 1260 cm^-1^. There is also the recognizable ∼1159 cm⁻¹ marker that confirms the NS1 type. In the pure NS1 sample, another weaker signal appears near 1230 cm^-1^; however, its strength is not anywhere close to that of the 1260 cm^-1^ band. When DENV-2 NS1 mixes with 10⁻² diluted HS, the 1260 cm^-1^ band stays strongly visible all the way down to 5 ng/mL. Even though its intensity falls off as we lower the NS1 concentration, the band sticks around. The raw data gets messy due to overlapping serum components. Nevertheless, the 1260 cm⁻¹ band appears as a distinct NS1-specific signal after deconvolution, indicating that we did not confuse some serum substance for NS1 activity. On the other hand, the weaker feature around 1230 cm⁻¹ is barely noticeable. For better comparison, if the model sticks to what worked for DENV-1, this small curve gets folded into a wider nearby band in all concentrations, and it makes a negligible contribution to the spectrum. Its peak position and peak area (∼0) are also not changing with concentration. In contrast, the 1260 cm^-1^ band supports the involvement of NS1’s vibrational modes; similar to DENV-1, the peak location varies very little, and the peak area declines with decreasing NS1 concentration as shown in Figure S13J and S13L. In conclusion, these observations show that the 1260 cm^-1^ band gives the strongest and most useful signal for spotting DENV-2 in HS. It differs from the usual ∼1159 cm^-1^ marker, which gets lost among other serum bands, because it stands out clearly after separating the data and can reliably indicate the presence of DENV-2 NS1 with the lowest concentration also.

In Figure S8, the DENV-3-specific diagnostic region—marked by the dark orange shading—is studied more using Gaussian peak deconvolution, as shown in Figure S14. This area matches where DENV-1 was checked first, denoted by light gray shading. Peak 1 at 954 cm^-1^ showcases the unique spectral marker for DENV-3 NS1. Figure S14A shows that the pure DENV-3 NS1 spectrum is mainly defined by a peak at around 954 cm^-1^. In the mixture of DENV-3 NS1 with HS diluted 10⁻², the spectrum changes due to vibrations from both NS1 and the serum. At 5 × 10^5^ ng/mL, the NS1 signal is super strong, and the 954 cm^-1^ peak is clearly visible. However, when the NS1 amount drops, other peaks from the serum start to take over. This makes the DENV-3 signal weaker. However, even when the NS1 concentration is lowest (5 ng/mL - see Figure S14H), the 954 cm^-1^ peak stays. Both raw data and the analyzed breakdown confirm this, showing that this feature keeps being visible despite lots of interference from the serum.

The concentration-dependent peak position and area are shown in Figures S14I and S14J. As the NS1 concentration goes down, the peak shifts to higher wavenumbers – meaning there are small changes in its local vibrational environment. At the same time, the peak area gets smaller when the NS1 concentration drops, showing that NS1 contributes less to the vibrations in the serum matrix. Besides the ∼954 cm^-1^ band, there is another band around ∼985 cm^-1^, from the strongest parts of the DENV-3 spectrum, and it mostly shows up at higher concentrations. It is much weaker at lower concentrations and doesn’t last as long as the 954 cm^-1^ peak, though. These results show that the ∼954–960 cm^-1^ band stays detectable throughout all concentrations and is the most reliable spectral marker for DENV-3 NS1 in HS. Also, its change depends on concentration, making it a strong fingerprint to identify the DENV-3 serotype in complex biological samples.

The DENV-4 NS1 has two important areas for diagnosis, marked in dark and light blue in Figure S8D. The deconvolution results for the first area are seen in Figure S15, while the second area shows up in Figure S16. These regions together give a special spectral fingerprint for DENV-4 and help tell it apart reliably, even when there is HS mixed in. For the first region, the main band is around 767 cm^-1^. In Figure S15A, the spectrum of pure DENV-4 NS1 fits well with two Gaussian curves. The big peak here is at around 767 cm⁻¹ (Peak 1, blue curve). When you combine DENV-4 NS1 with diluted 10^-2^ HS, the spectral profile stays pretty much the same as the pure NS1 at concentrations of 1 × 10^5^ and 5 × 10^4^ ng/mL (Figures S15B and S15C), showing the NS1 signals are still very strong. At 5 × 10^3^ ng/mL, though, we start seeing the serum’s effects. However, even down to really low concentrations—from 500 to 5 ng/mL—that distinctive 767 cm⁻¹ component stands out clearly in both the raw and deconvoluted spectra. In the HS control spectrum, the relevant feature is small compared to mixture and pure dengue samples. Analyzing the peak position and area in Figures S15I and S15J, the peak moves towards lower wavenumbers as NS1 concentration drops. Similarly, the peak area also decreases with concentration, showing less influence from NS1-derived vibrations in the serum. Therefore, this band appears across all concentrations, making it a great sensitive diagnostic marker for DENV-4 NS1. The second diagnostic area, seen in Figure S16, shows the typical DENV-4 marker at around ∼1065 cm^-1^. Figure S16A demonstrates that the pure DENV-4 NS1 spectrum can be broken down into four parts, with Peak 3 (highest intensity peak, blue curve) matching the ∼1065 cm^-1^ band. The spectra for mixtures at 5 × 10^5^ ng/mL appear very much like the pure NS1 spectrum, suggesting that NS1 vibrations take over here too. As the NS1 concentration drops, the serum starts making its presence felt more; however, the big hump at ∼1065 cm^-1^ stays obvious throughout— down to 5 ng/mL. The analysis of the fitted peak parameters shows that the position of the ∼1065 cm^-1^ band stays essentially constant, which means excellent spectral stability. However, the integrated peak area gets smaller as the NS1 concentration goes down, showing less protein presence. The broad feature that persists in comparison to the HS control indicates that it originates from NS1 and highlights its value as a serotype-specific identifier.

Together, the deconvolution analysis shows that having both the ∼767 and ∼1065 cm⁻¹ bands at once creates a unique fingerprint for DENV-4 NS1. Both features were present throughout the tested concentration range and stand out from the HS background, suggesting these bands can reliably identify the DENV-4 serotype in complicated biological samples. Additionally, having two separate markers makes identifying the DENV-4 serotype way more confident and reliable than counting on just one spectral feature.

In conclusion, the results of the deconvolution analysis of all four dengue NS1 serotypes at 50 and 5 ng/mL in an HS (10^-2^) are shown in Figure 4. In spite of the heavy background from serum, the distinct bands characteristic of each serotype are clearly visible in their distinct spectra region, with the lowest concentration also; indicating the efficiency of the deconvolution analysis method on the SERS platform.

**Figure 4.**
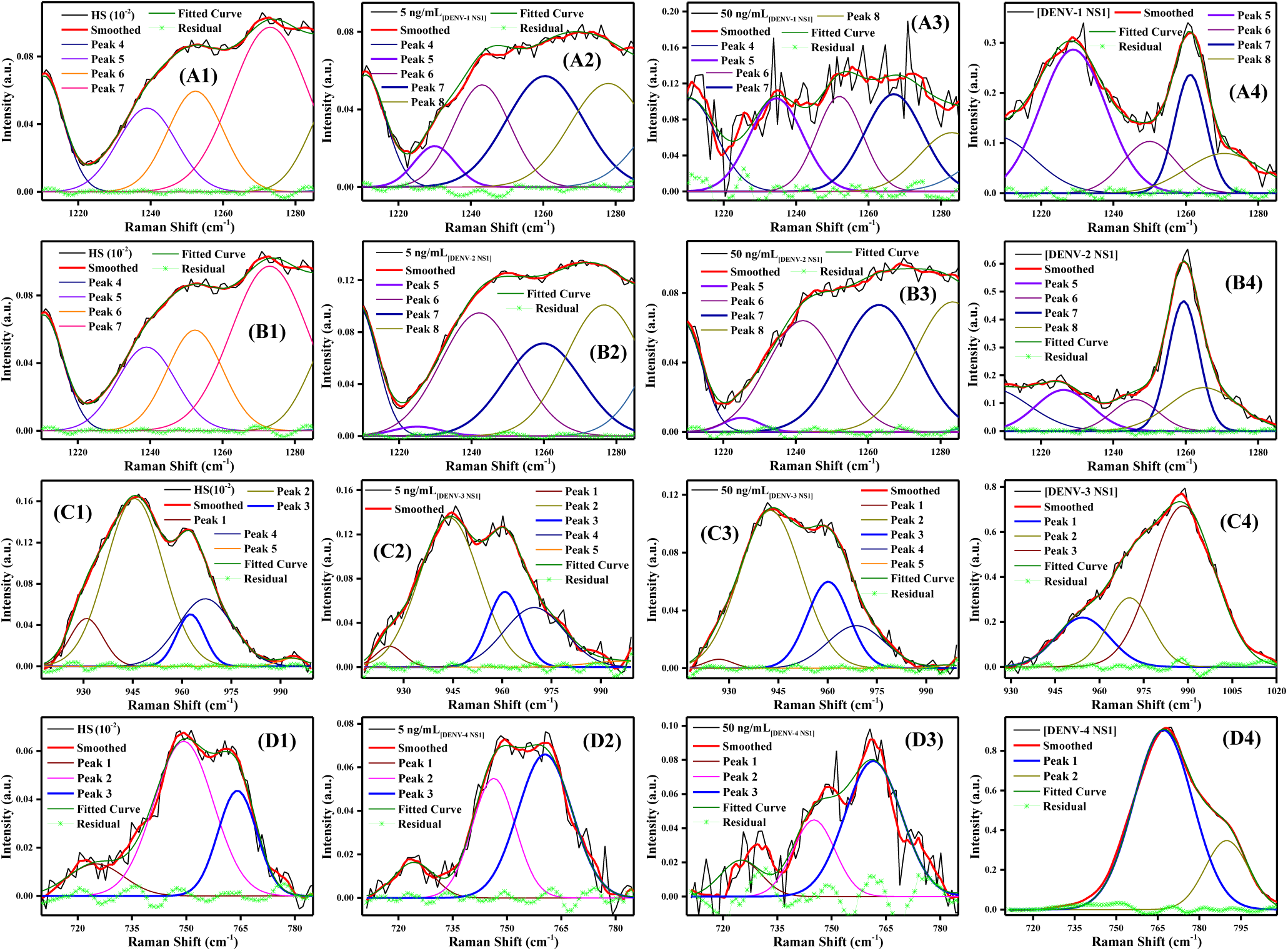
Deconvolution of Raman spectra: (A1–A4) DENV-1 NS1, (B1–B4) DENV-2 NS1, (C1–C4) DENV-3 NS1, and (D1–D4) DENV-4 NS1. Series 1–4 represent the diluted HS (10⁻²), mixtures of corresponding NS1 (5 and 50 ng/mL) with HS (10⁻²), and pure serotype-specific NS1, respectively.

These results demonstrate that the variations in the amino acid sequence of the NS1 proteins from the four dengue types are reflected in their SERS fingerprints and can be interpreted using spectral deconvolution, even in the presence of a complex serum mixture. Even at concentrations as low as 5 ng/mL, the method is still able to detect those crucial features. This demonstrates the method’s sensitivity and importance, which makes it ideal for identifying the serotype NS1 in practical samples. This study establishes a link between real-world clinical use and lab-based SERS testing. It’s significant to be able to determine whether a person has dengue and the type of infection using a single, quick test. It outperforms previous techniques because it enables quick scanning and precise strain identification. Additionally, this procedure is excellent for point-of-care applications because of its low prices, minimum preparation requirements, and highly repeatable outcomes. This approach has the potential to significantly improve the efficacy and accessibility of dengue diagnosis. These results provide a foundation for the development of rapid, affordable, and portable SERS diagnostic instruments for dengue. They might also be useful for other illnesses, where identifying and differentiating strains is crucial for infection monitoring and patient care.

## CONCLUSION

The present study demonstrates the ability of SERS to distinguish and detect serotype-specific NS1 proteins among DENV serotypes by using signature peaks in vibrational Raman spectra of NS1 proteins. Based on the early and accurate identification of NS1 proteins from DENV serotypes, our findings show that SERS holds great potential as a powerful tool for enhancing the accuracy of DENV serotype infection diagnosis. The marker peaks for every DENV serotype are evidently visible in the spectra of all serotype NS1 proteins. Additionally *in-vitro* solutions of human sera with varying concentration of NS1 proteins from all serotypes demonstrate the marker peaks in serotypes, thus distinguishing between serotypes and identifying sequence variation in NS1 proteins irrespective of their biological source. This demonstrates that SERS is a promising method with good sensitivity, fast analysis, minimal sample preparation requirements, and accurate early identification of serotype-specific DENV infections. This approach has the potential to improve disease control and management strategies and deepen our understanding of DV pathogenesis.

## ACKNOWLEDGEMENT

The authors would like to acknowledge Indian Council of Medical Research (ICMR) and Council of Scientific and Industrial Research (CSIR) for support of core instrumental facilities. One of the authors M.G has been supported by a Senior Research Fellowship Grant from the University Grants Commission. The authors would like to thank all staff of the core facilities in providing technical support.

