## Supplementry File for "Serotype-Specific Detection of Non-Structural Protein 1 from Dengue Viruses by Surface-Enhanced Raman Spectroscopy: An Enhanced Precision Diagnosis"

### Supporting Information

#### Surface-Enhanced Raman Spectroscopy-Based Detection of Nonstructural-1 (NS1) from Dengue Virus Serotypes: An Enhanced Diagnostic Precision

No. of Pages—**18**

No. of Figures— **16**

No. of Table— **01**

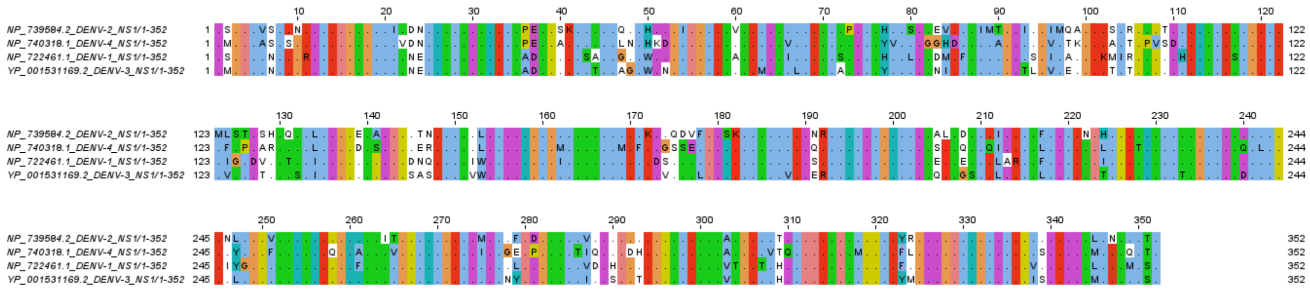

**Figure S1.** Multiple sequence alignment of full-length NS1 proteins from different DENV serotypes, highlighting amino acid residue variations. Residues are represented in single-letter code, with only non-conserved (dissimilar) residues shown.

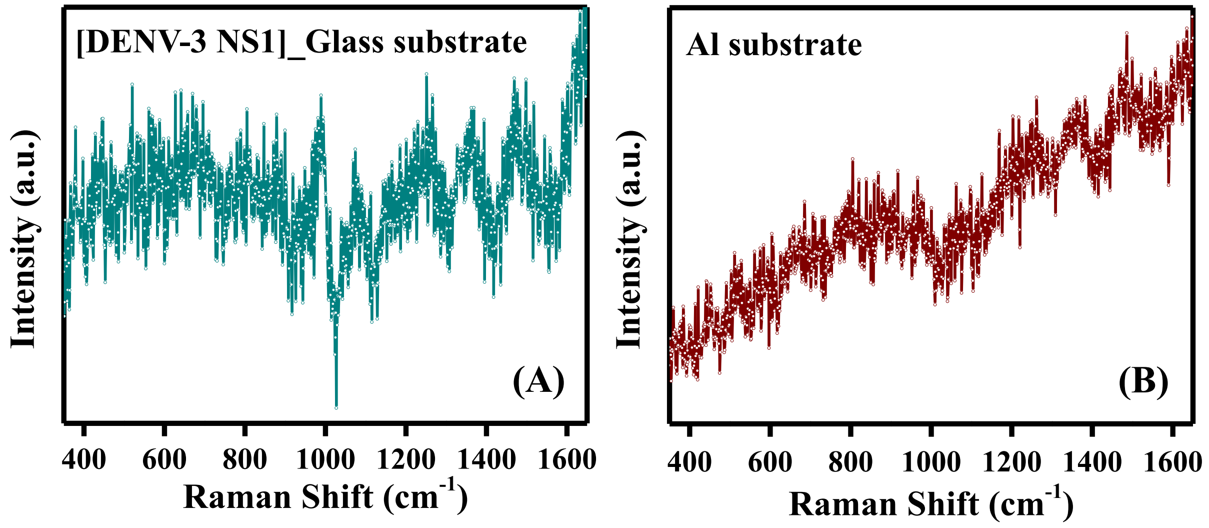


**Figure S2. (**A) Raman spectrum of DENV-3 NS1 acquired on a glass substrate. (B) Raman spectrum of aluminum foil employed as the SERS substrate.

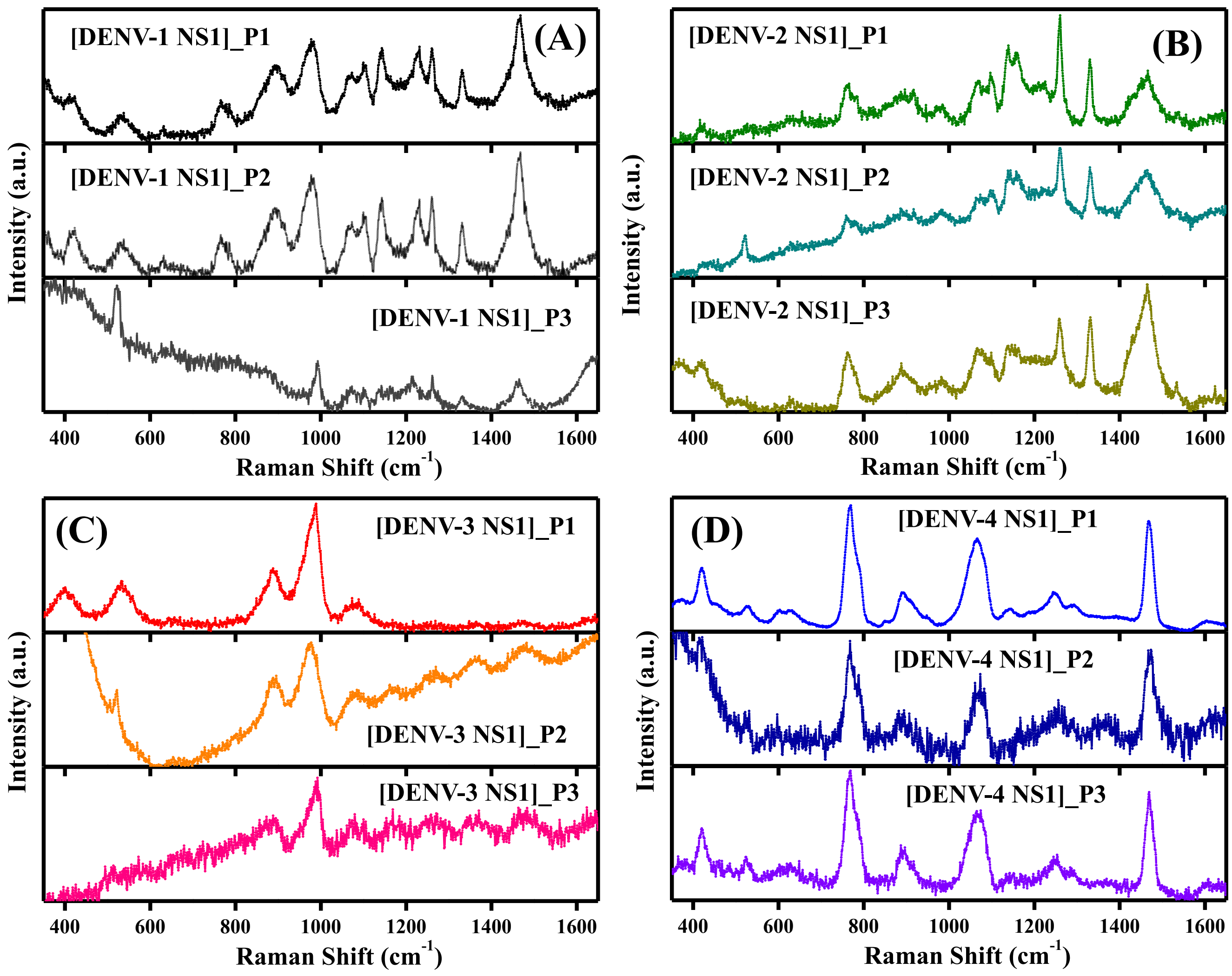


**Figure S3.** SERS spectra of purified recombinant NS1 proteins from different DENV serotypes (1.0 μg), exhibiting distinct spectral fingerprints. (A) DENV-1 NS1, (B) DENV-2 NS1, (C) DENV-3 NS1, and (D) DENV-4 NS1. Spectra were acquired from three different positions (P1, P2, and P3) on the same sample for each serotype.

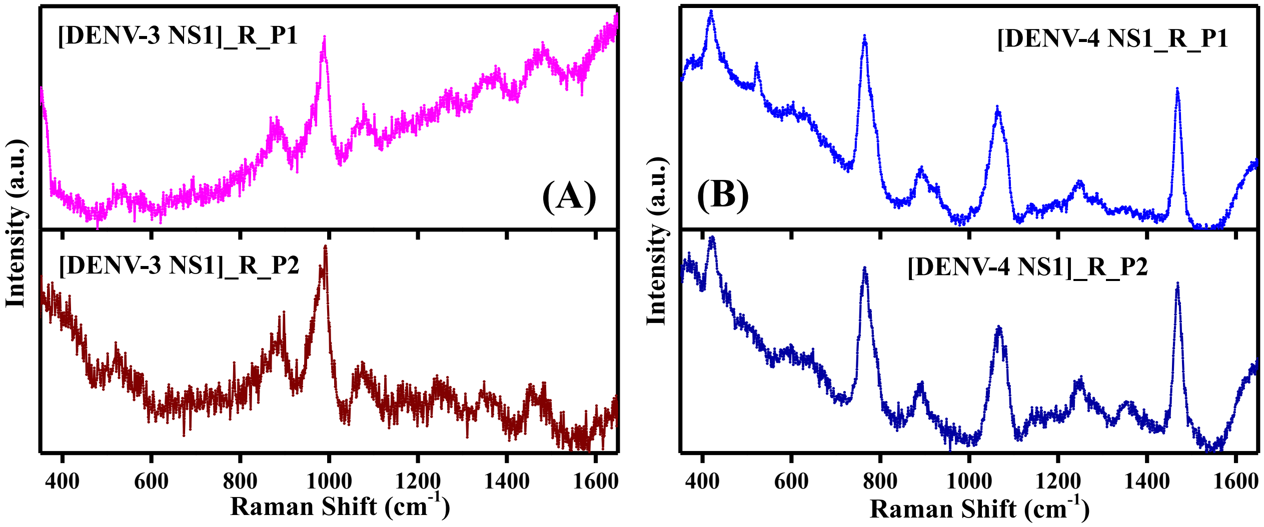


**Figure S4.** Reproducibility assessment of SERS spectra: (A) DENV-3 NS1 (R) and (B) DENV-4 NS1 (R), demonstrating spectral consistency across measurements. Spectra were acquired from two different positions (P1 and P2) on the same sample for each serotype.

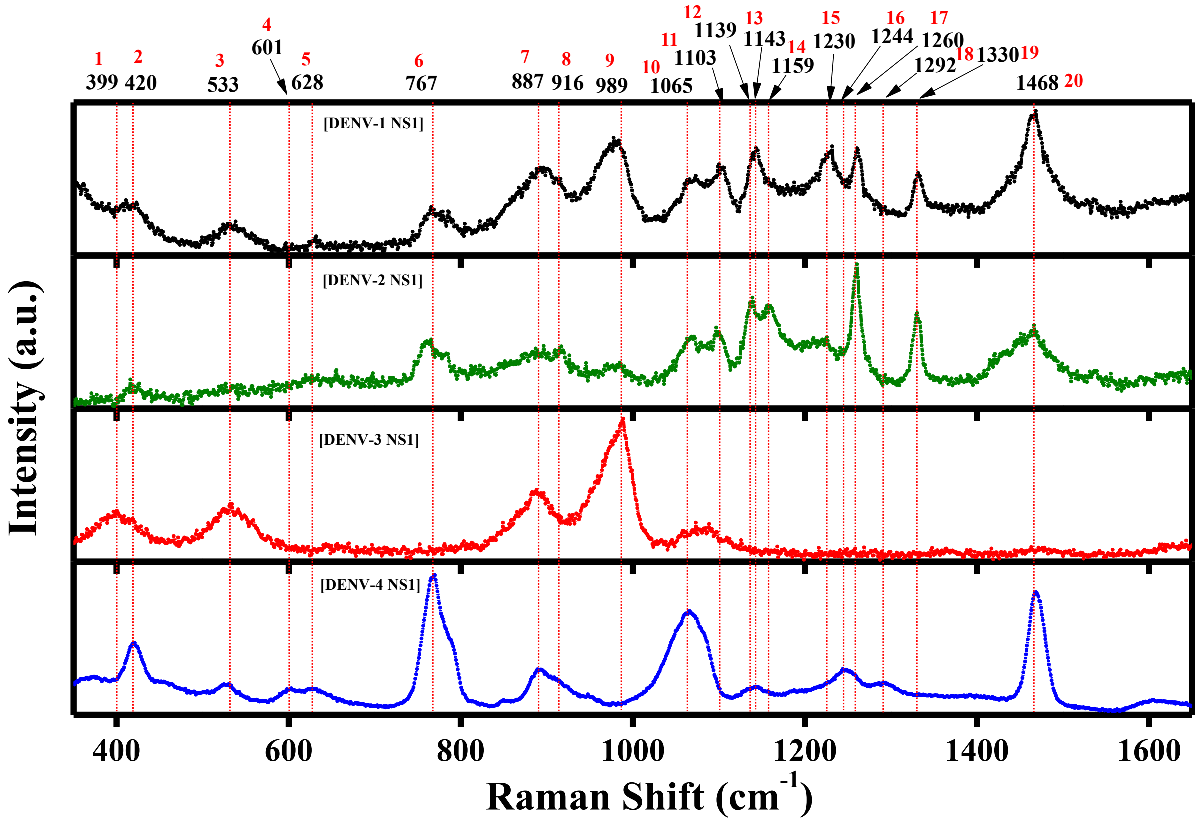


**Figure S5.** SERS spectra of purified recombinant NS1 proteins from different DENV serotypes (1.0 μg), exhibiting distinct spectral fingerprints. Lines indicate peak positions, with corresponding Raman shift values (black) and peak numbers labeled (red).

**Table S1.** Tabulated SERS peaks of NS1 proteins from different DENV serotypes, showing distinct spectral fingerprints. Color intensity reflects peak intensity (darker = higher), and corresponding amino acid assignments are included.

| Peak Position (cm$^{-1}$) / Stereotype | 399 | 420 | 533 | 601 | 628 | 767 | 887 | 916 | 989 | 1065 | 1103 | 1139 | 1143 | 1159 | 1230 | 1244 | 1260 | 1292 | 1330 | 1468 |
|---|---|---|---|---|---|---|---|---|---|---|---|---|---|---|---|---|---|---|---|---|
| [DENV-1 NS1] | A | P | P | A | P | P | P | M | P | P (D) | P(D) | M | P | A | P | A | P | A | P | P |
| [DENV-2 NS1] | A | P | A | A | P | P | P | P | P | P (D) | P(D) | P(D) | M | P(D) | P(B) | A | P | A | P | P |
| [DENV-3 NS1] | P | A | P | A | A | A | P | A | P | M (B) | A | A | A | A | A | A | A | A | A | P |
| [DENV-4 NS1] | A | P | P | P | P | P | P | M | A | P | A | P | P | A | A | P | A | P | A | P |

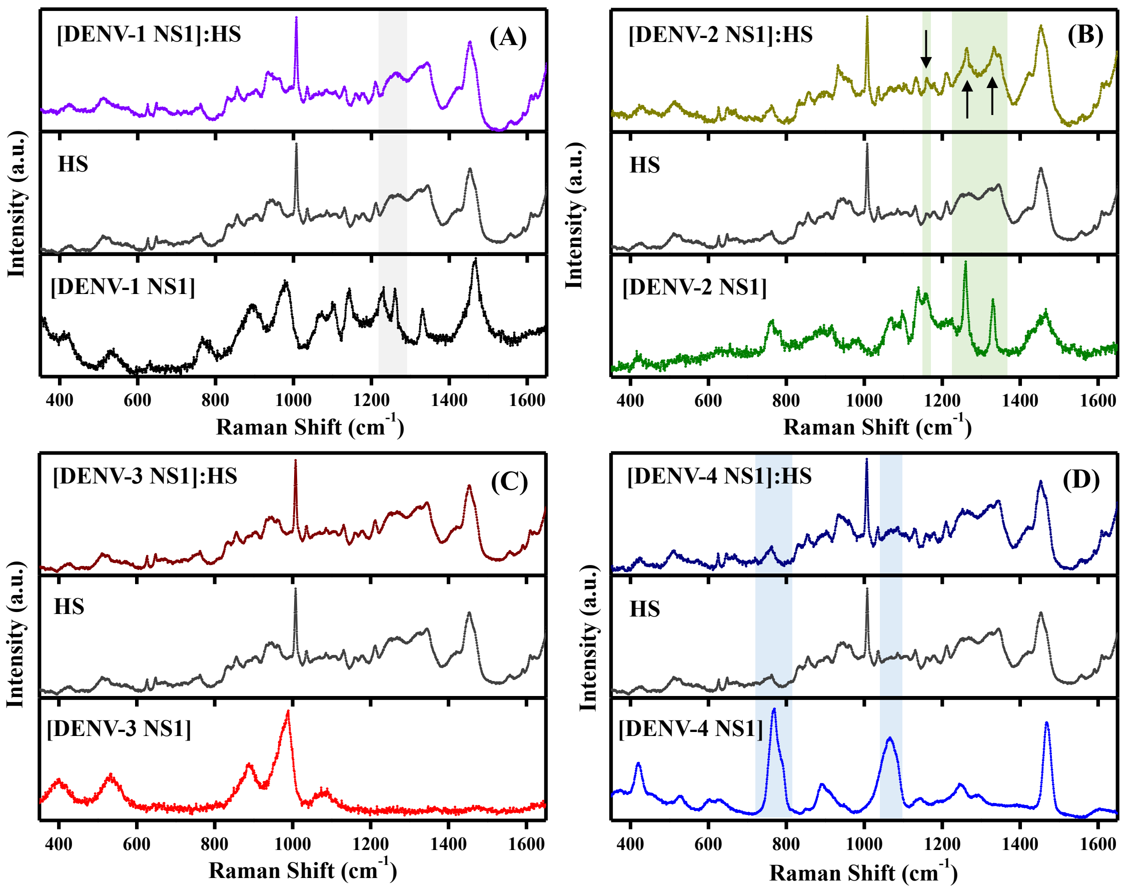


**Figure S6.** (A–D) SERS spectra of NS1 proteins from different DENV serotypes, human serum (HS), and their 1:1 mixture. Shaded regions denote characteristic signature peaks of each NS1 protein, which are weakly discernible for (A) DENV-1 and (D) DENV-4, most prominent for (B) DENV-2, and not clearly observable for (C) DENV-3 in the mixture.

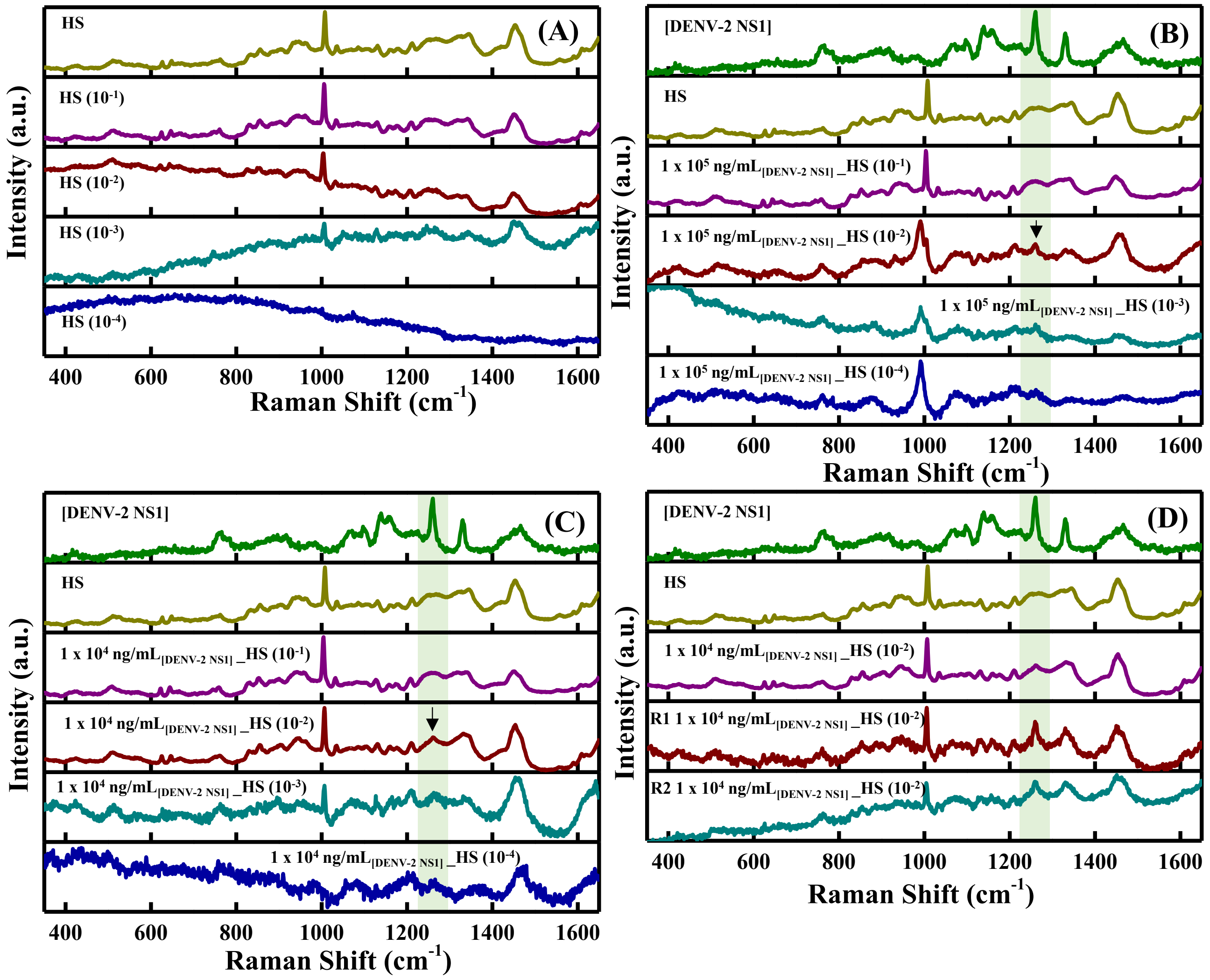


**Figure S7.** SERS spectra: (A) Human serum (HS), pure and diluted ($10^{-1}$–$10^{-4}$). (B) Pure DENV-2 NS1, HS, and DENV-2 NS1 (1 x $10^5$ ng/mL) mixed 1:1 with diluted HS ($10^{-1}$–$10^{-4}$). (C) Pure DENV-2 NS1, HS, and DENV-2 NS1 (1 x $10^4$ ng/mL) mixed 1:1 with diluted HS ($10^{-1}$–$10^{-4}$). (D) Reproducibility assessment (n = 3) of SERS spectra for 10 μg/mL DENV-2 NS1 with HS ($10^{-2}$). Shaded regions denote characteristic signature peaks.

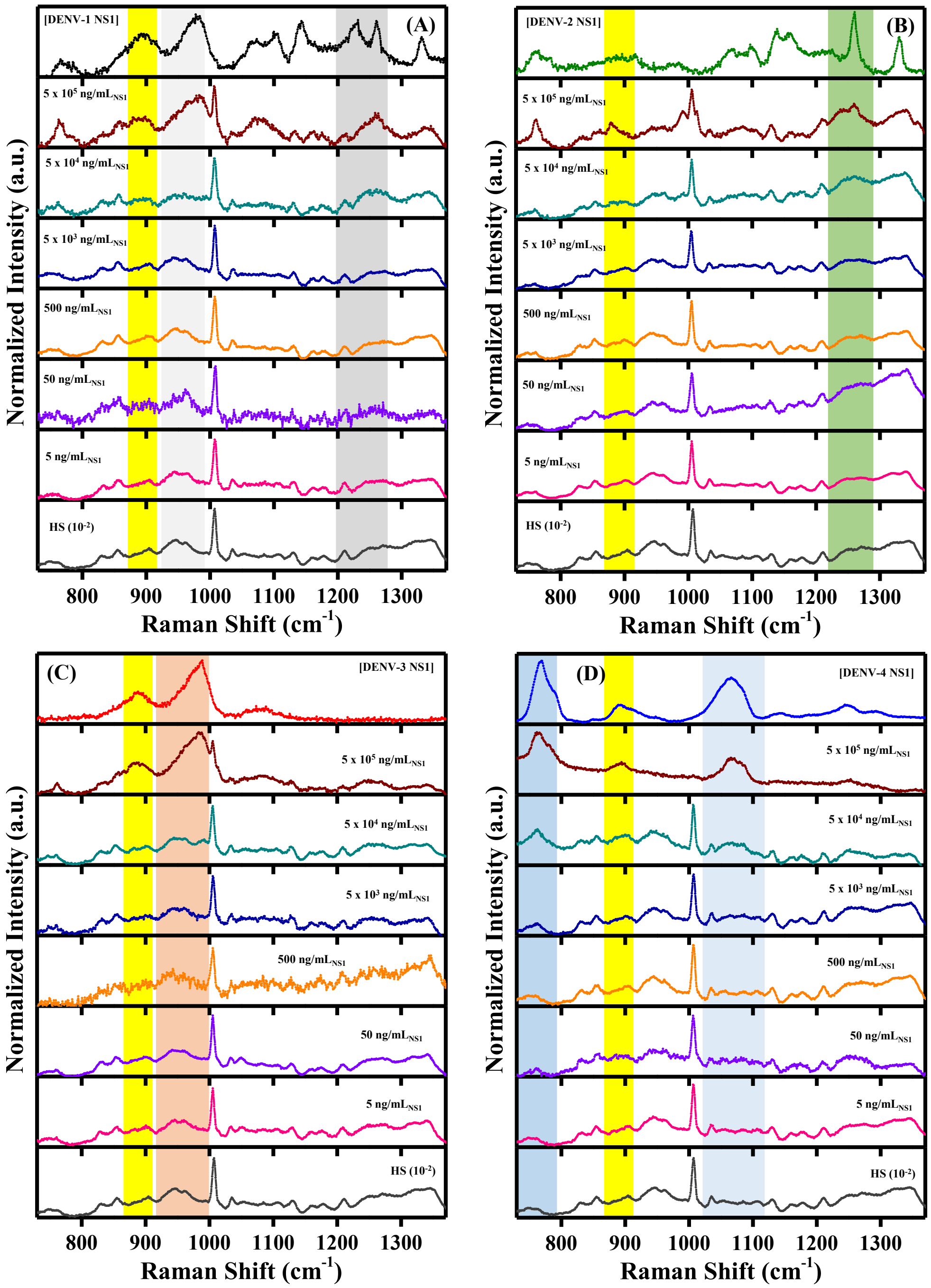


**Figure S8.** SERS spectra: (A) DENV-1 NS1, (B) DENV-2 NS1, (C) DENV-3 NS1, and (D) DENV-4 NS1, each showing pure NS1, mixtures of NS1 (5 x 10$^5$–5 ng/mL) with HS (10$^{-2}$), and HS (10$^{-2}$). Shaded regions denote characteristic signature peaks: yellow indicates common peak regions; dark grey and light grey represent characteristic and primary serotype-specific peaks for DENV-1 NS1, respectively; dark green for DENV-2 NS1; orange for DENV-3 NS1; and dark blue and light blue represent primary and characteristic serotype-specific peaks for DENV-4 NS1, respectively.

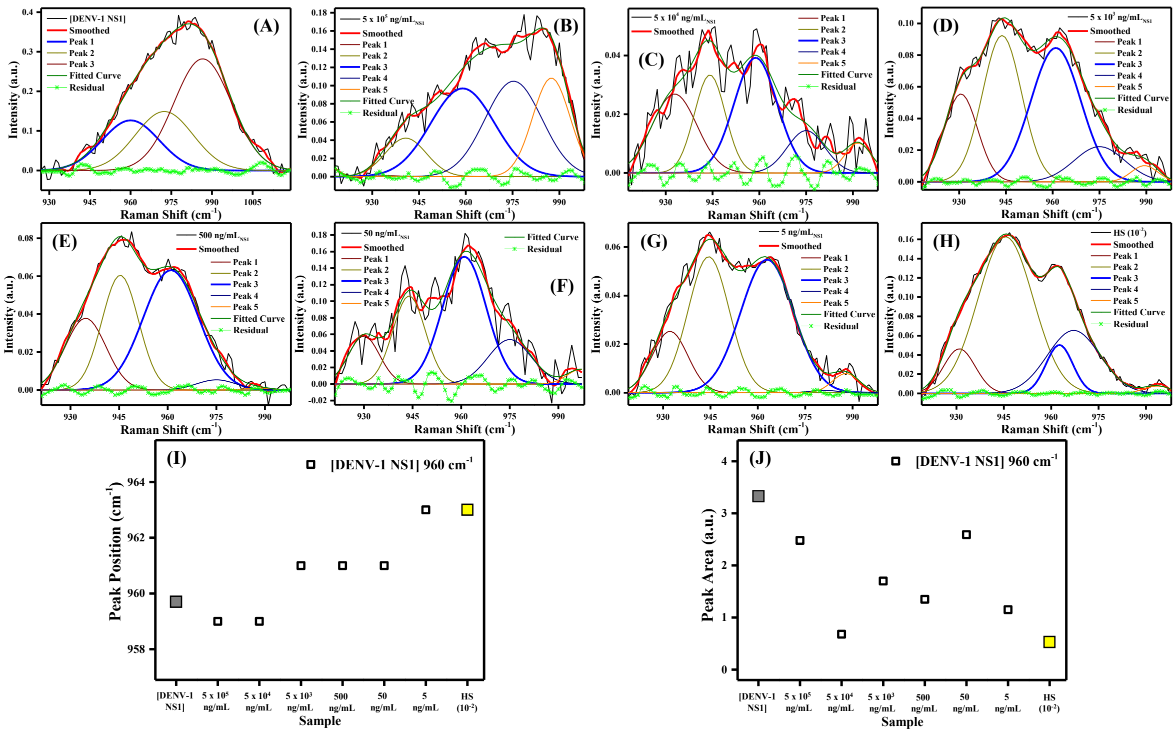

**Figure S9.** Deconvolution of Raman spectra of DENV-1 NS1: (A) Pure DENV-1 NS1; (B–G) mixtures of NS1 (5 x $10^5$, 5 x $10^4$, 5 x $10^3$, 500, 50, and 5 ng/mL) with HS ($10^{-2}$); and (H) HS ($10^{-2}$). (I) Peak position and (J) peak area as a function of concentration for the ~960 $cm^{-1}$ (square) bands. Filled grey symbols denote pure NS1, while the filled yellow symbol represents HS ($10^{-2}$).

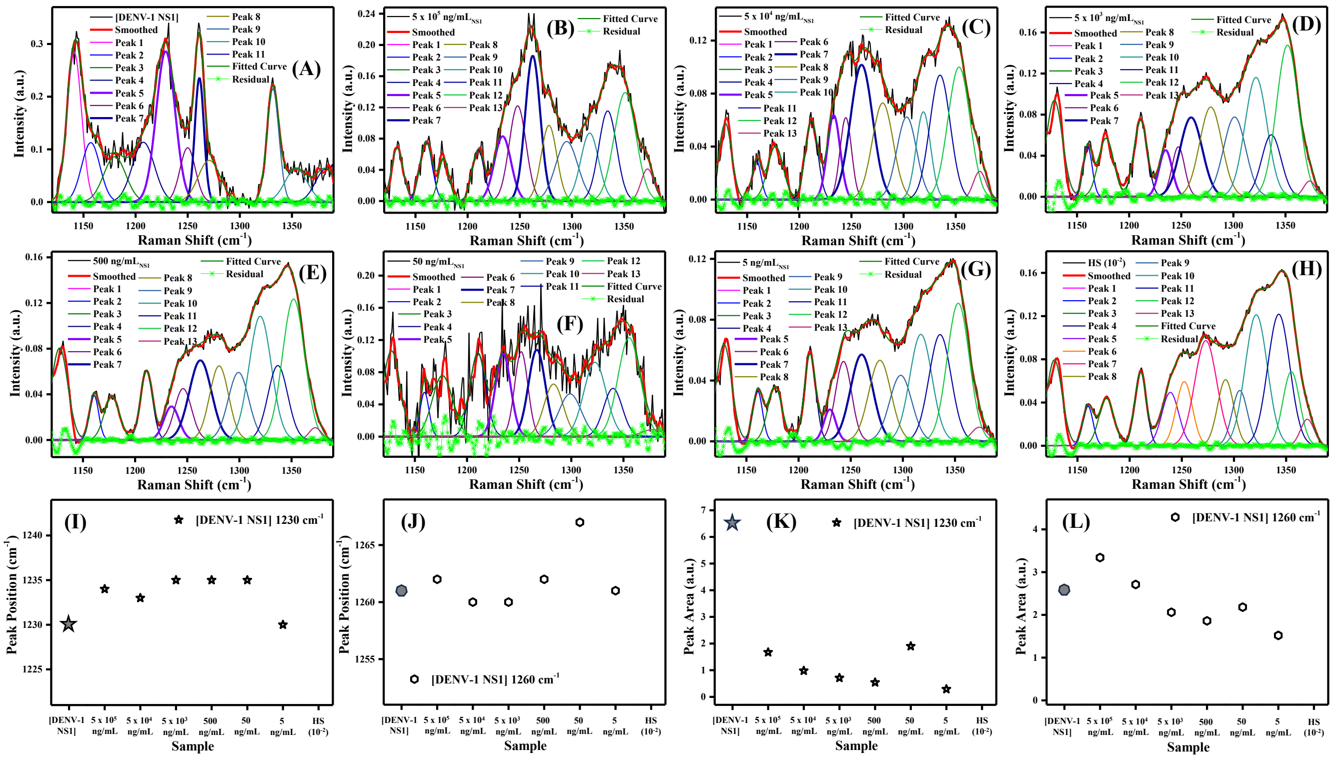

**Figure S10.** Deconvolution of Raman spectra of DENV-1 NS1: (A) Pure DENV-1 NS1; (B–G) mixtures of NS1 (5 x $10^5$, 5 x $10^4$, 5 x $10^3$, 500, 50, and 5 ng/mL) with HS ($10^{-2}$); and (H) HS ($10^{-2}$). (I–J) Peak position and (K–L) peak area as a function of concentration for the ~1230 $cm^{-1}$ (star) and ~1260 $cm^{-1}$ (circle) bands. Filled grey symbols denote pure NS1.

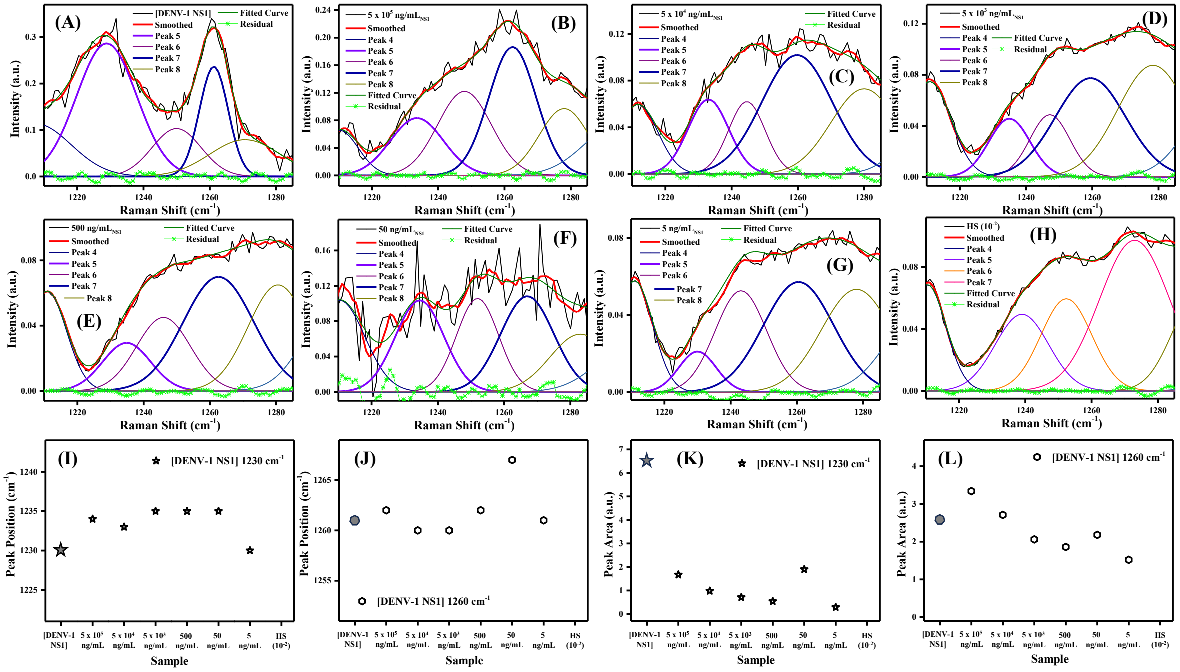

**Figure S11.** Zoomed-in view of deconvolution of Raman spectra of DENV-1 NS1: (A) Pure DENV-1 NS1; (B–G) mixtures of NS1 (5 x $10^5$, 5 x $10^4$, 5 x $10^3$, 500, 50, and 5 ng/mL) with HS ($10^{-2}$); and (H) HS ($10^{-2}$). (I–J) Peak position and (K–L) peak area as a function of concentration for the ~1230 $cm^{-1}$ (star) and ~1260 $cm^{-1}$ (circle) bands. Filled grey symbols denote pure NS1.

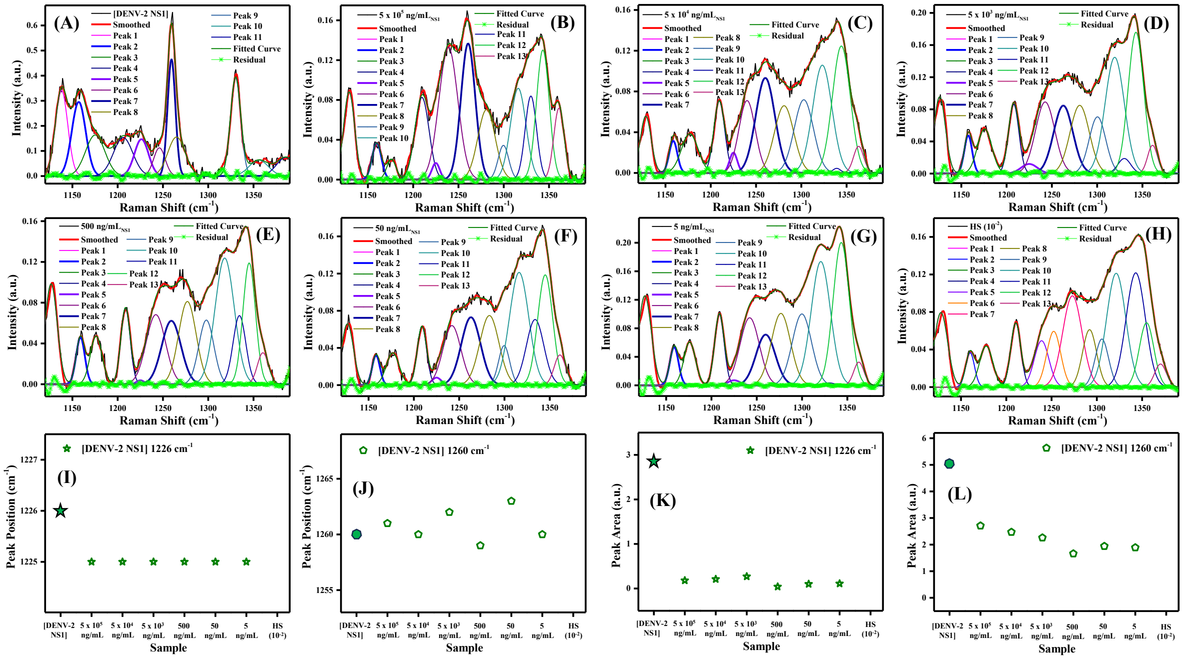

**Figure S12.** Deconvolution of Raman spectra of DENV-2 NS1: (A) Pure DENV-2 NS1; (B–G) mixtures of NS1 (5 x $10^5$, 5 x $10^4$, 5 x $10^3$, 500, 50, and 5 ng/mL) with HS ($10^{-2}$); and (H) HS ($10^{-2}$). (I–J) Peak position and (K–L) peak area as a function of concentration for the ~1230 $cm^{-1}$ (star) and ~1260 $cm^{-1}$ (circle) bands. Filled green symbols denote pure DENV-2 NS1.

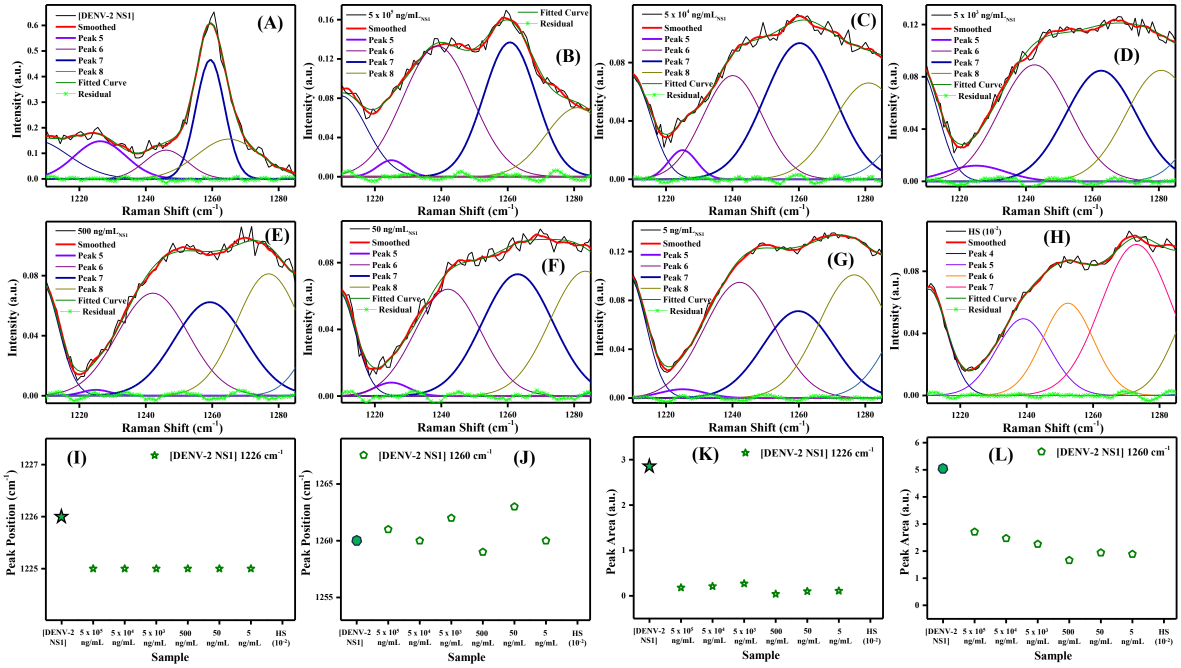


**Figure S13.** Zoomed-in view of deconvolution of Raman spectra of DENV-2 NS1: (A) Pure DENV-2 NS1; (B–G) mixtures of NS1 (5 x $10^5$, 5 x $10^4$, 5 x $10^3$, 500, 50, and 5 ng/mL) with HS ($10^{-2}$); and (H) HS ($10^{-2}$). (I–J) Peak position and (K–L) peak area as a function of concentration for the ~1230 $cm^{-1}$ (star) and ~1260 $cm^{-1}$ (circle) bands. Filled grey symbols denote pure NS1.

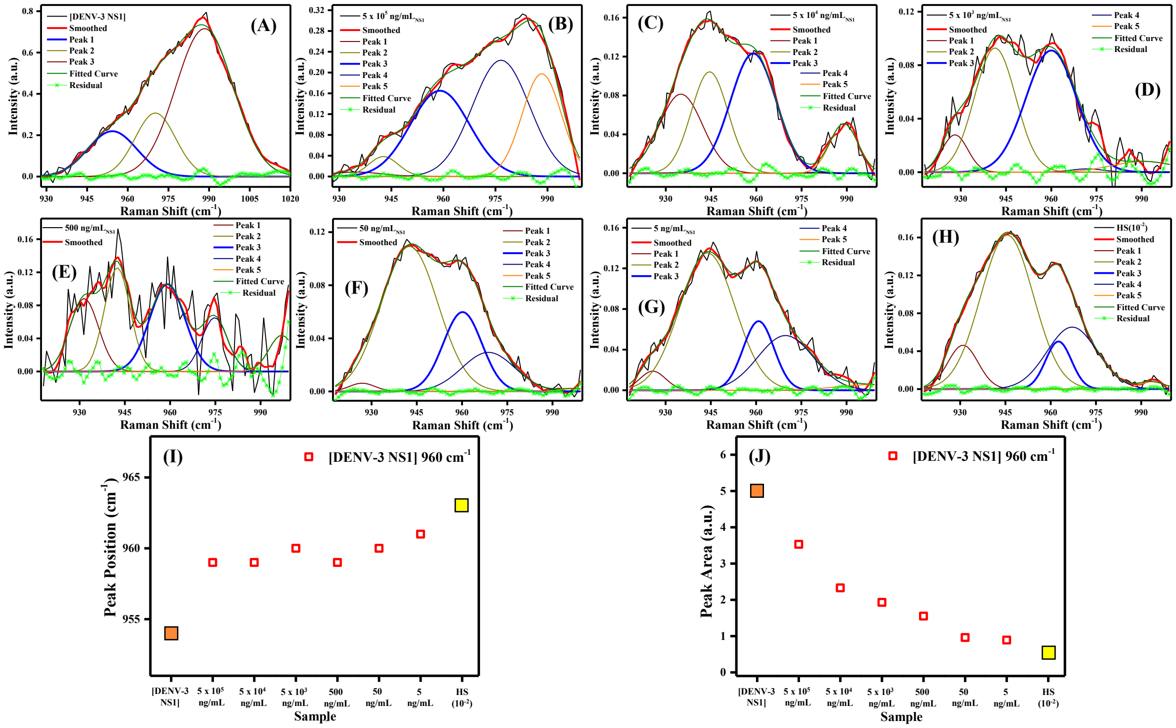


**Figure S14.** Deconvolution of Raman spectra of DENV-3 NS1: (A) Pure DENV-3 NS1; (B–G) mixtures of NS1 ($5 \times 10^5$, $5 \times 10^4$, $5 \times 10^3$, 500, 50, and 5 ng/mL) with HS ($10^{-2}$); and (H) HS ($10^{-2}$). (I) Peak position and (J) peak area as a function of concentration for the ~960 $cm^{-1}$ (square) bands. Filled orange symbols denote pure NS1, while the filled yellow symbol represents HS ($10^{-2}$).

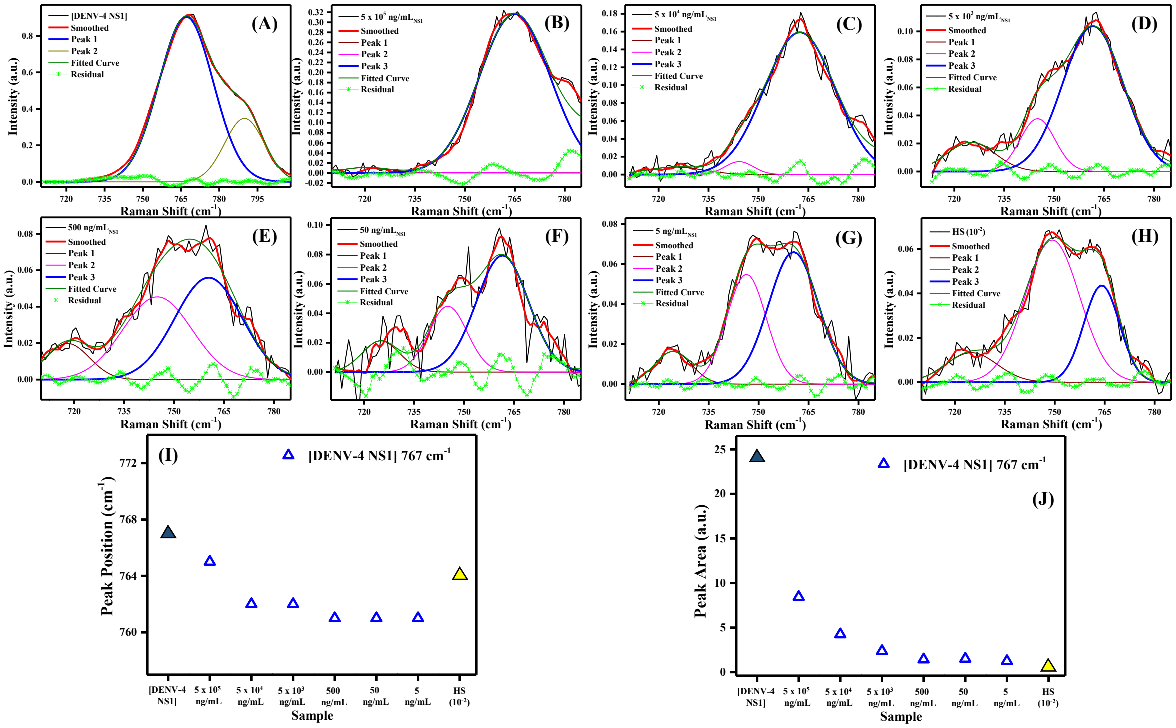


**Figure S15.** Deconvolution of Raman spectra of DENV-4 NS1: (A) Pure DENV-4 NS1; (B–G) mixtures of NS1 ($5 \times 10^5$, $5 \times 10^4$, $5 \times 10^3$, 500, 50, and 5 ng/mL) with HS ($10^{-2}$); and (H) HS ($10^{-2}$). (I) Peak position and (J) peak area as a function of concentration for the ~767 $cm^{-1}$ (triangle) bands. Filled blue symbols denote pure NS1, while the filled yellow symbol represents HS ($10^{-2}$).

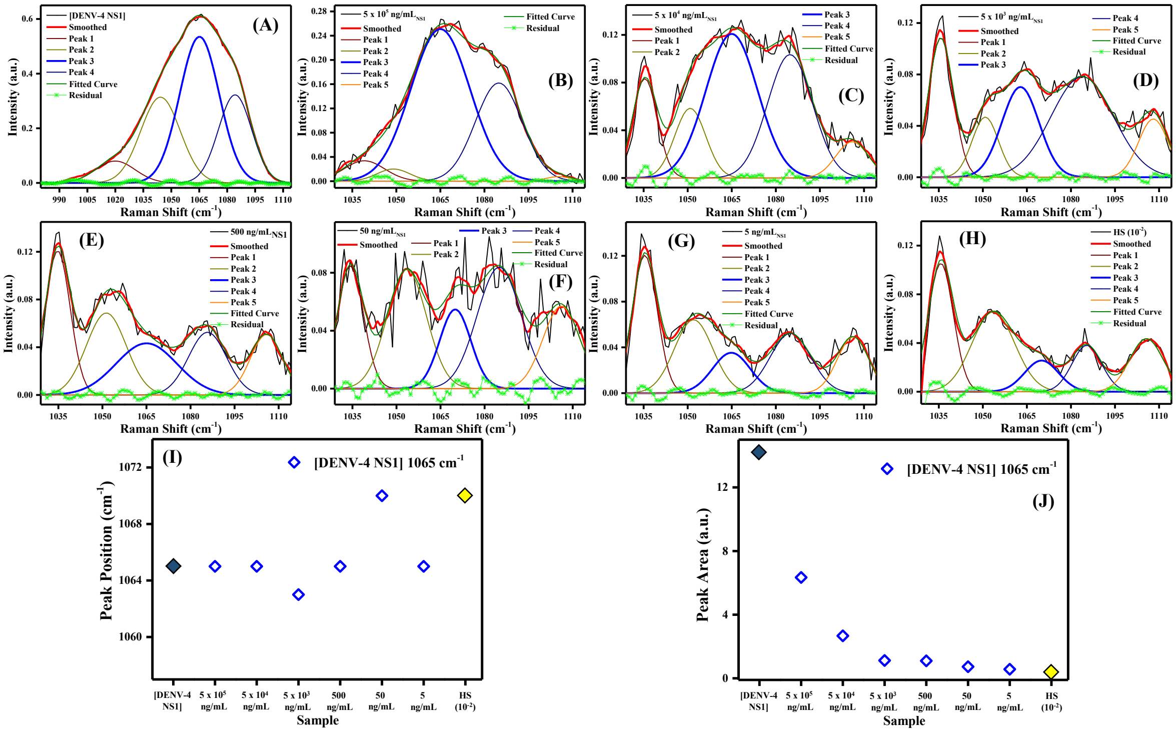

**Figure S16.** Deconvolution of Raman spectra of DENV-4 NS1: (A) Pure DENV-4 NS1; (B–G) mixtures of NS1 (5 x $10^5$, 5 x $10^4$, 5 x $10^3$, 500, 50, and 5 ng/mL) with HS ($10^{-2}$); and (H) HS ($10^{-2}$). (I) Peak position and (J) peak area as a function of concentration for the ~1065 $cm^{-1}$ (diamond) bands. Filled blue symbols denote pure DENV-4 NS1, while the filled yellow symbol represents HS ($10^{-2}$).
